# Experience-dependent information routing through the basolateral amygdala

**DOI:** 10.1101/2023.08.02.551710

**Authors:** Pantelis Antonoudiou, Bradly Stone, Phillip L.W. Colmers, Aidan Evans-Strong, Najah Walton, Grant Weiss, Jamie Maguire

## Abstract

The basolateral amygdala (BLA) is an emotional processing hub and is well-established to influence both positive and negative valence processing. Selective engagement of a heterogeneous cell population in the BLA is thought to contribute to this flexibility in valence processing. However, how this process is impacted by previous experiences which influence valence processing is unknown. Here we demonstrate that previous positive (EE) or negative (chronic unpredictable stress) experiences differentially influence the activity of specific populations of BLA principal neurons projecting to either the nucleus accumbens core or bed nucleus of the stria terminalis. Using chemogenetic manipulation of these projection-specific neurons we can mimic or occlude the effects of chronic unpredictable stress or enriched environment on valence processing to bidirectionally control avoidance behaviors and stress-induced helplessness. These data demonstrate that previous experiences influence the responsiveness of projection-specific BLA principal neurons, biasing information routing through the BLA, to govern valence processing.

## Introduction

The ability to rapidly evaluate contexts or stimuli and determine whether the experience is positive or negative is highly adaptive and essential for survival. Valence processing is a dynamic process which influences the determination of whether something is considered “good” or “bad”. Positive and negative valence processing are often thought of as opposing processes mediated by unique circuits (for review see (Correia and Goosens, 2016)). However, it has also been suggested that valence processing is governed by a “mixed selectivity” model in which individual neurons may be recruited by different stimuli under different contexts and can thereby contribute to disparate functions in different contexts (for review see (Coley et al., 2021)). This concept of flexible valence processing is difficult to reconcile with the “Divergent Paths” circuit motif which is thought to underlie valence switching in brain regions such as the amygdala (for review see (Tye, 2018)), involving divergent information routing to distinct downstream targets.

The basolateral amygdala (BLA) is a critical hub for both positive and negative valence processing which is thought to involve divergent downstream circuits (for review see (Daviu et al., 2019)). For example, single unit recording demonstrated that projection-specific neurons are differentially activated by reward prediction versus a prediction of an aversive stimulus (Beyeler et al., 2016) (for review see (Beyeler et al., 2018)). Optogenetic stimulation of BLA neurons projecting to the nucleus accumbens (BLA-NAc) supports reward seeking (Namburi et al., 2015); whereas, optogenetic stimulation of BLA neurons projecting to the central amygdala (BLA-CeA) induces place avoidance (Namburi et al., 2015). Similarly, activation of BLA neurons projecting to the ventral hippocampus (BLA-vHPC) induces avoidance behaviors (Felix-Ortiz et al., 2013). Thus, it is clear that distinct, projection-specific principal neurons in the BLA participate in positive and negative valence processing (for review see (Daviu et al., 2019)). Thus, we envision that valence processing is a dynamic process involving the recruitment of specific ensembles and downstream circuits.

Deficits in valence processing have been associated with numerous psychiatric illnesses, including anxiety, depression, and post-traumatic stress disorder (PTSD). For example, attentional bias for negatively valanced stimuli has been demonstrated in individuals with anxiety disorders (MacLeod et al., 2019), with negative interpretations of ambiguous sentences (Richards and French, 1992), scenarios (Hirsch and Mathews, 1997), and surprised faces (Park et al., 2016). (for review see (Daviu et al., 2019)). Attentional bias and the tendency to attend to negative stimuli has also been shown in patients with depression (Mennen et al., 2019; Peckham et al., 2010) and PTSD (Bryant and Harvey, 1997; Iacoviello et al., 2014). These findings suggest that valence processing is compromised in association with mood disorders.

Chronic stress is a major risk factor for psychiatric illnesses (Mazure, 1998; Tennant, 2002). Previous trauma exposure increases attentional bias towards a new threat (Blekić et al., 2021) and experimentally, chronic stress in animal models has been shown to alter valence behaviors, increasing negative valence processing (Monteiro et al., 2015; Willner et al., 1987) (for review see (Tran and Gellner, 2023; Willner, 1984, 1997, 2017; Willner et al., 1992)). The circuits implicated in valence process overlap with those implicated in stress (for review see (Daviu et al., 2019)). Therefore, we propose that previous stress experience may influence the recruitment of valence-specific neurons and circuits. Conversely, enriched environment has been shown to alter valence processing in mice, decreasing negative valence processing (Lin et al., 2011). Therefore, we propose that previous experiences, both positive and negative, may influence the recruitment of neurons and circuits underlying valence processing.

Here we demonstrate that chronic unpredictable stress (CUS) alters the electrophysiological properties of principal BLA neurons in a projection-specific manner, facilitating the activation of BnST-projecting BLA principal neurons (BLA-BnST) and suppressing the activity of NAc core-projecting BLA (BLA-NAcc) principal neurons. CUS exposure also alters the stress-induced activation and synchrony within subsets of BLA principal neurons. Conversely, enriched environment (EE) alters the electrophysiological properties of BLA principal neurons such that the activity of BLA-NAcc neurons is enhanced, and BLA-BnST neurons is suppressed. EE also influences the stress-induced activation and synchrony within subsets of BLA principal neurons in opposition to that of CUS. Chemogenetic activation of BLA-BnST and suppression of BLA-NAcc is sufficient to mimic the behavioral consequences of CUS; whereas, chemogenetic activation of BLA-NAcc and suppression of BLA-BnST can prevent the increased negative valence processing following CUS. Conversely, chemogenetic activation of BLA-NAcc and suppression of BLA-BnST mimics the behavioral effects of EE; whereas chemogenetic activation of BLA-BnST and suppression of BLA-NAcc is sufficient to prevent the reduced negative valence processing following EE. These data demonstrate that previous positive and negative experiences bias information routing through the BLA to facilitate information flow through specific downstream circuits to reduce or increase negative valence processing, respectively. These findings provide a cellular and circuit mechanism mediating the impact of life experiences on the risk for psychiatric illnesses.

## Results

### Chronic unpredictable stress alters the electrophysiological properties of BLA principal neurons

In order to characterize the effects of chronic unpredictable stress (CUS) on the excitability of principal neurons in the BLA, we performed whole-cell patch clamp recordings in current clamp mode and measured the intrinsic membrane properties and generated input-output curves in control (CNT) and CUS mice. The recording schematic and action potential waveforms are shown in Figure 1A. Following chronic stress, BLA principal neurons exhibit an increase in excitability, significantly firing more, in response to lower input currents compared to CNT mice (Figure 1B). For example, the action potential frequency in response to a 30pA current injection is significantly increased in BLA principal neurons from mice subjected to CUS (8.50 ± 1.82 Hz) compared to CNT (0.80 ± 0.57 Hz) (Figure 1B). We compared the differences across extracted electrophysiological properties including action potential waveforms between BLA principal neurons from CNT and CUS mice (Supplemental Figures 1-2). An increase in the variability of numerous electrophysiological properties of BLA principal neurons was observed in BLA principal neurons following CUS (Supplemental Figure 2b). We noticed from the distribution plot of the resting membrane potential (RMP) in CNT and CUS animals (Figure 1C) that the variability in the RMP of cells from CUS mice can be separated into two putative populations based on the mode of the control cells: one hyperpolarized and one depolarized compared to CNT (Figure 1D) which can be further visualized when plotting the rheobase across the RMP (Figure 1E). These findings indicate that CUS may differentially modulate the excitability of distinct BLA principal cell populations.

**Figure 1:**
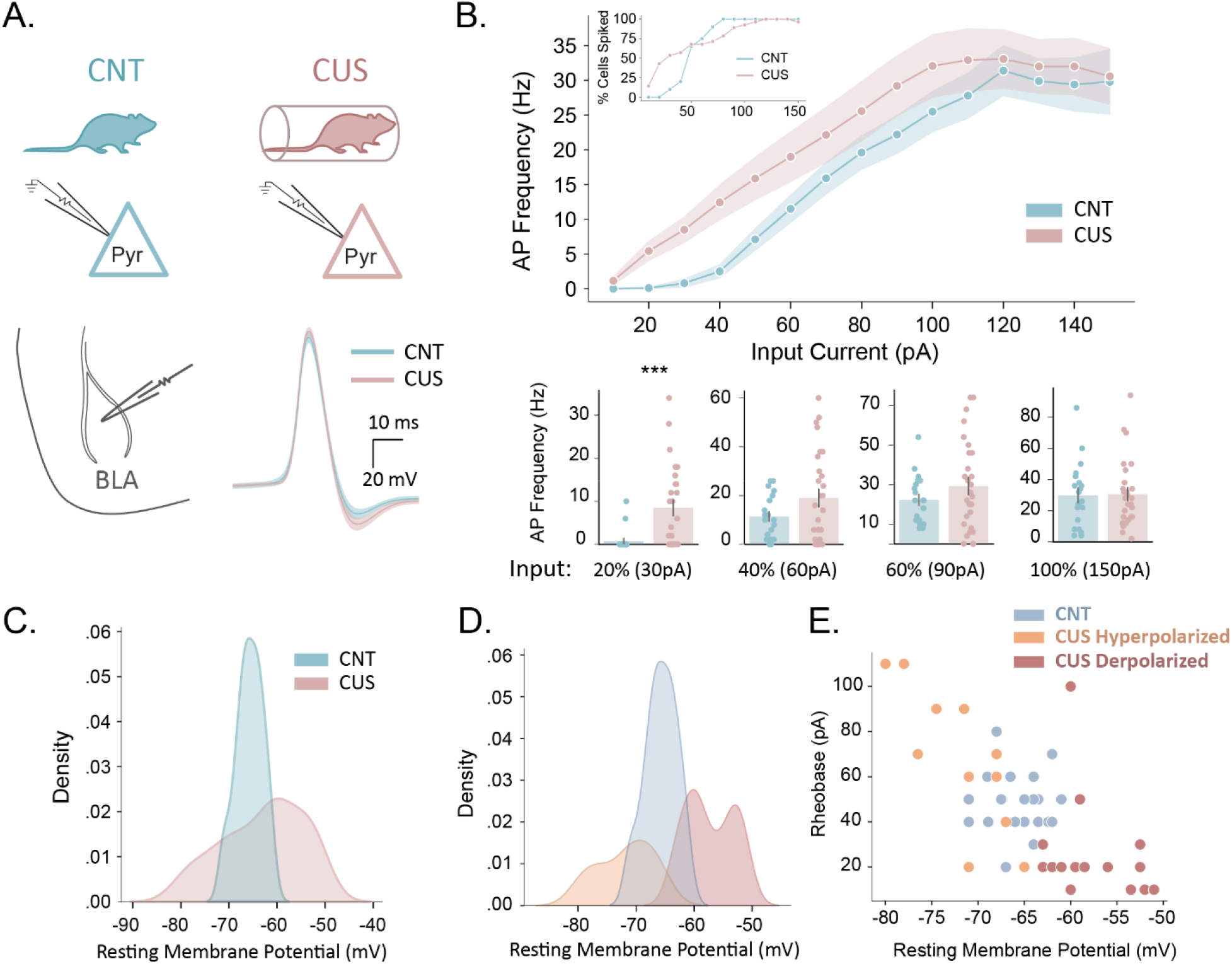
CUS alters electrophysiological properties of BLA principal neurons. A) Schematic diagram illustrating BLA principal cells recorded from CNT and CUS mice along with the average waveform of recorded cells. B) Input-output curve (top), along with summary bar plots (bottom) showing that CUS increases the firing rate of principal cells at low input thresholds. C) The variability of the resting membrane potential (RMP) is higher in CUS compared to CNT principal cells. D) Distribution plot and E) scatterplot illustrating putative depolarized and hyperpolarized clusters resulting from splitting CUS cells based on the mode of CNT RMP and assessed by a permutation-based MANOVA (Stats Table 1).

**Figure 2:**
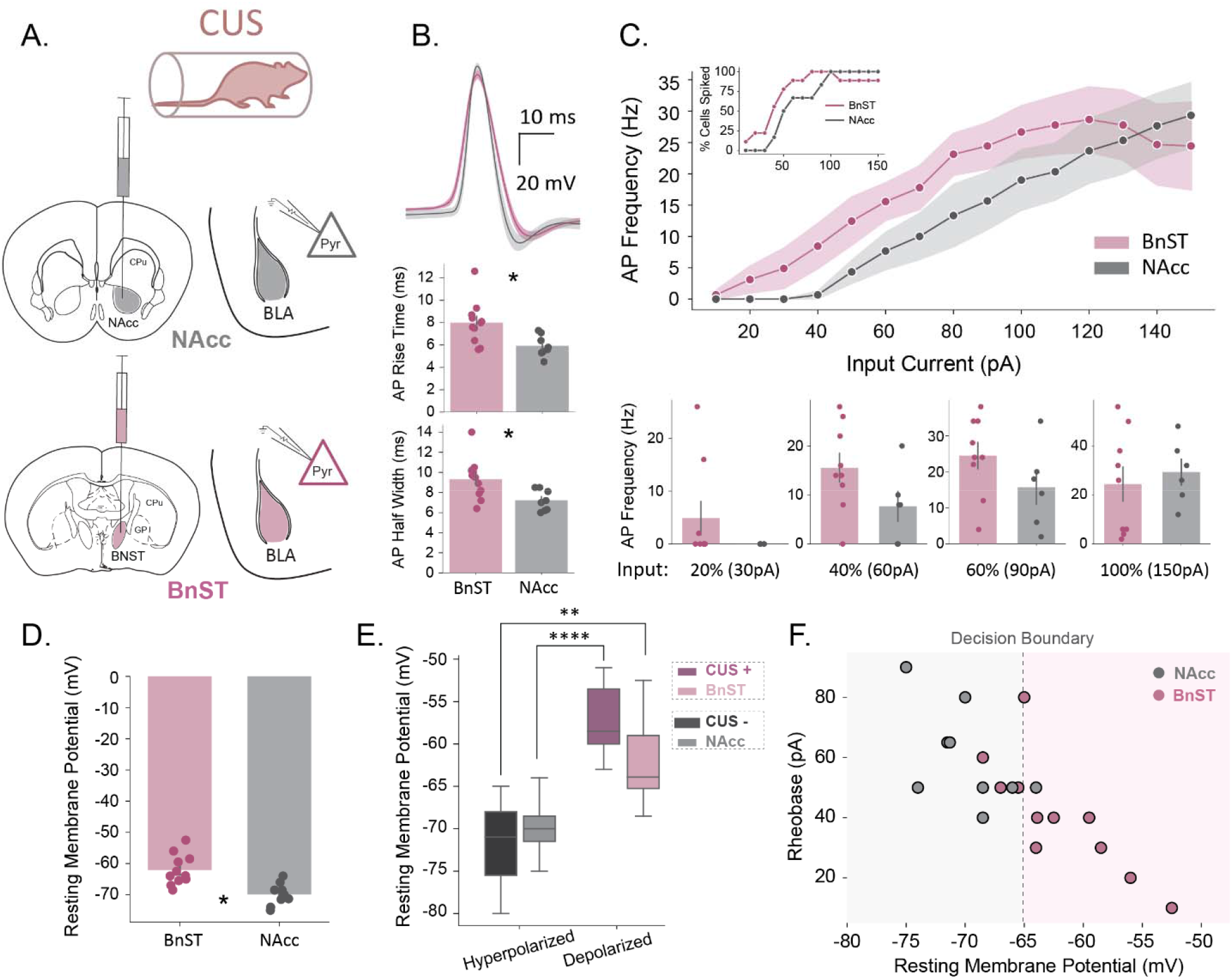
Projection-specific impact of CUS on BLA principal neurons. **A)** Schematic diagram illustrating the targeting and recording of BLA principal cells projecting to either NAcc or BnST. **B)** Average action potential waveform (top) along with quantifications of rise-time (middle) and half-width (bottom). **C)** Input-output curve (top), along with summary bar plots (bottom). **D)** Resting membrane potential of BnST and NAcc cells. **E)** Comparison of resting membrane potential of NAcc and BnST cells with CUS hyperpolarized and depolarized clusters. **F)** Scatter plot of rheobase against resting membrane potential decision boundary of a gaussian naïve bayes classifier trained on CUS depolarized and hyperpolarized clusters (Supplemental Table 1, row 0).

### Chronic unpredictable stress alters the electrophysiological properties of BLA principal neurons in a projection-specific manner

To determine whether the increased variability in the electrophysiological properties of BLA principal neurons following chronic stress is due to differential effects on distinct populations of projection neurons, we used retrograde tracing to identify projection-specific populations and determine the impact of CUS on their electrophysiological properties (Figure 2A). We found that the action potential (AP) rise time and half width were reduced in BLA principal neurons projecting to the NAcc (BLA-NAcc) (rise time: 5.93 ± 0.34 ms; half width: 7.22 ± 0.37 ms) compared to those projecting to the BnST (BLA-BnST) (rise time: 7.99 ± 0.58 ms; half width: 9.32 ± 0.61 ms) (Figure 2B) with no significant difference in the input-output curves (Figure 2C). Comparisons in the electrophysiological properties between BLA-BnST and BLA-NAcc neurons from CUS mice are shown in Supplemental Figure 3. Remarkably, we find a significant difference in the RMP in select BLA principal neuron populations following CUS with BLA-NAcc neurons being more hyperpolarized (-69.87 ± 1.19 mV) compared to BLA-BnST (-62.08 ± 1.48 mV) (Figure 2D). Following CUS, the BLA-NAcc neurons are not significantly different from the CUS hyperpolarized population (-71.86 mV) and BLA-BnST neurons are not significantly different from the CUS depolarized population (-57.24 ± 1.00 mV); Conversely, both BLA-NAcc and BLA-BnST neurons are significantly different from CUS depolarized and hyperpolarized populations, respectively (Figure 2E). Furthermore, a linear classifier showed that the best predictor of BLA-BnST and BLA-NAcc populations after CUS was the separation of CUS cells by the mode of the RMP in control cells used here (Figure 2F; Supplemental Table 1). Overall, these results support that CUS induces changes in the electrophysiological properties of BLA principal neurons to promote information flow from BLA to BnST and inhibit information flow to the NAcc.

### BLA-BnST activation and BLA-NAcc suppression mimics the behavioral effects of chronic unpredictable stress

Our results suggest that CUS may preferentially facilitate information flow from the BLA to the BnST over NAcc projections. To causally test whether enhancement of BLA-BnST pathway and suppression of BLA-NAcc pathway is sufficient to induce behavioral deficits similar to CUS we expressed GqM3 in BLA-BnST cells and GiM4 in BLA-NAcc cells (CNT BnST+NAcc-) and tested their behavior alongside CNT and CUS mice (Figure 3A-B). We did not find any significant differences in the distance traveled, time spent, and entries to center in the open field (OF) between the 3 groups (Figure 3C). However, we observed a reduction in the total distance traveled and an increase in the total time immobile in CUS and CNT BnST+NAcc-compared to CNT mice (Supplemental Figure 3A), suggesting that CUS and CNT BnST+ NAcc-mice exhibit increased freezing behavior indicative of the behavioral expression of fear. CUS and CNT BnST+NAcc-mice also exhibited increased avoidance behaviors, evident from the reduced distance traveled, time spent, and entries in the light zone of the light dark box (LD) compared to CNT mice (Figure 3D). Moreover, CUS and CNT BnST+NAcc-mice had lower latency to immobility and reduced total time immobile in the tail suspension test (TST) compared to CNT mice (Figure 3E). Therefore, simultaneous activation of BLA-BnST and inactivation of BLA-NAcc projections seems to be sufficient to induce CUS-like behavioral impairments in C57 mice. To further test whether suppressing the BLA-BnST and activating the BLA-NAcc projections can prevent the behavioral impairments of CUS, we also tested the behavior of mice that underwent CUS and expressed GiM4 in BLA-BnST cells and GqM3 in BLA-NAcc cells (CUS BnST-NAcc+). We found that CUS BnST-NAcc+ mice had closer behavioral profiles to CNT mice than CUS mice as indicated by decreased avoidance behaviors (Figure 3 C-D), decreased stress-induced helplessness (Figure 3E), and unsupervised clustering of the behavioral outputs (Figure 3F). Specifically, all CUS and CNT BnST+NAcc-were in cluster 2 whereas all CNT along with 73.3 % of CUS BnST-NAcc+ mice fell into cluster 1 (Figure 3F). Thus, BLA-BnST suppression and BLA-Knack activation can reduce CUS-induced behavioral impairments.

**Figure 3:**
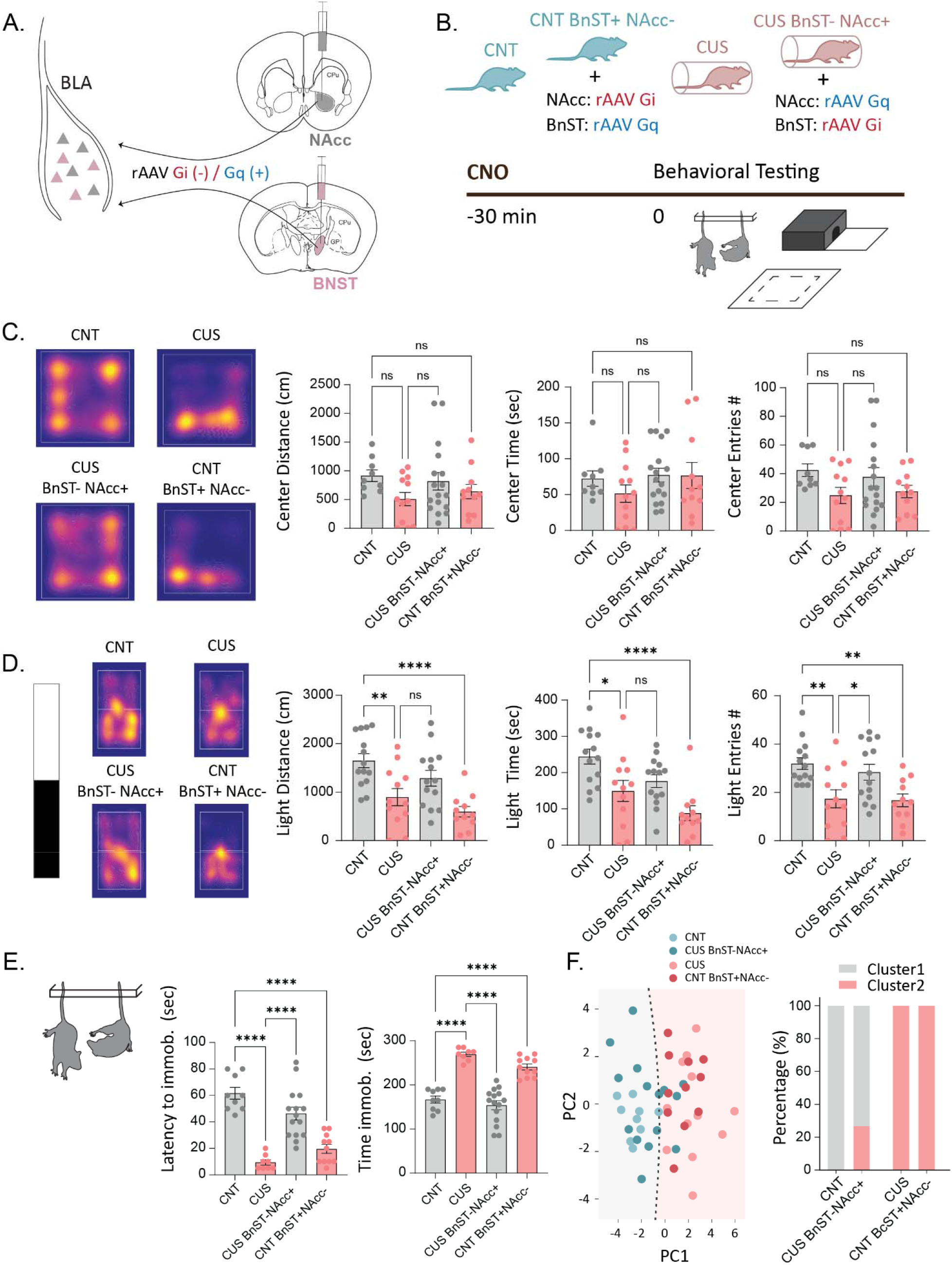
Activation of BLA-BnST pathway and inactivation of BLA-NAcc pathway mimics the behavioral effects of CUS. **A)** Schematic diagram illustrating targeting of BLA-BnST and BLA-NAcc pathway using retroAAV Gq/Gi DREADDs. **B)** Schematic diagram indicating the treatment groups (control: CNT, control with rAAV Gq in BnST and rAAV Gi in NAcc: CNT BnST-NAcc+, chronic unpredictable stress: CUS, chronic unpredictable stress with rAAV Gi in BnST and rAAV Gq in NAcc: CUS BnST+NAcc-), and that mice were injected with CNO (dose) 30 minutes prior to each behavioral test. **C)** Open field test representative heatmaps (left) and summary bar plots of distance, time, and entries in center zone across groups (right). **D)** Light Dark box test representative heatmaps (left) and summary bar plots of distance, time, and entries in light zone across groups (right). **E)** Tail suspension test summary bar plots quantifying latency to immobility and total time immobile. **F)** Clustering of the groups into two clusters using a GMM and the first three principal components. Decision boundary for the first 2 principal components (left) and percentage of each

### Enriched environment alters the electrophysiological properties of BLA principal neurons

The lack of variability in the electrophysiological properties of BLA principal neurons under control conditions raised the question about whether the standard homecage environment provided a lack of experience in valence processing leading to homogeneous electrophysiological properties. To test this hypothesis, we examined the impact of EE on the electrophysiological properties of BLA principal neurons (Figure 4A). Enriched environment increased the average action potential duration compared to CNT, increasing the peak to trough (16.38 ± 0.73 ms) and the AP half width (8.07 ± 0.67 ms) compared to CNT (peak to trough: 13.50 ± 0.64 ms; half width: 7.07 ± 0.49 ms) (Figure 4B). Enriched environment also altered the input-output curves in BLA principal neurons, reducing the frequency of action potentials fired in response to larger current injections (150 pA) (12.21 ± 2.27 Hz) compared to CNT (29.80 ± 4.66 Hz) (Figure 4C) and decreasing the input resistance of EE BLA principal neurons (227.41 ± 34.29 MΩ) compared to CNT (425.93 ± 40.26 MΩ) (Figure 4D). Similar to CUS, we observed an increase in the RMP in mice experiencing EE compared to CNT (Figure 4E, Supplemental Figure 5). The variability in the RMP in BLA principal neurons in mice housed under EE conditions can be separated into two putative clusters based on the mode of the RMP in control cells: one hyperpolarized and one depolarized compared to CNT (Figure 4F). These findings raise the possibility that EE differentially alters excitability of distinct BLA principal cell subpopulations.

**Figure 4:**
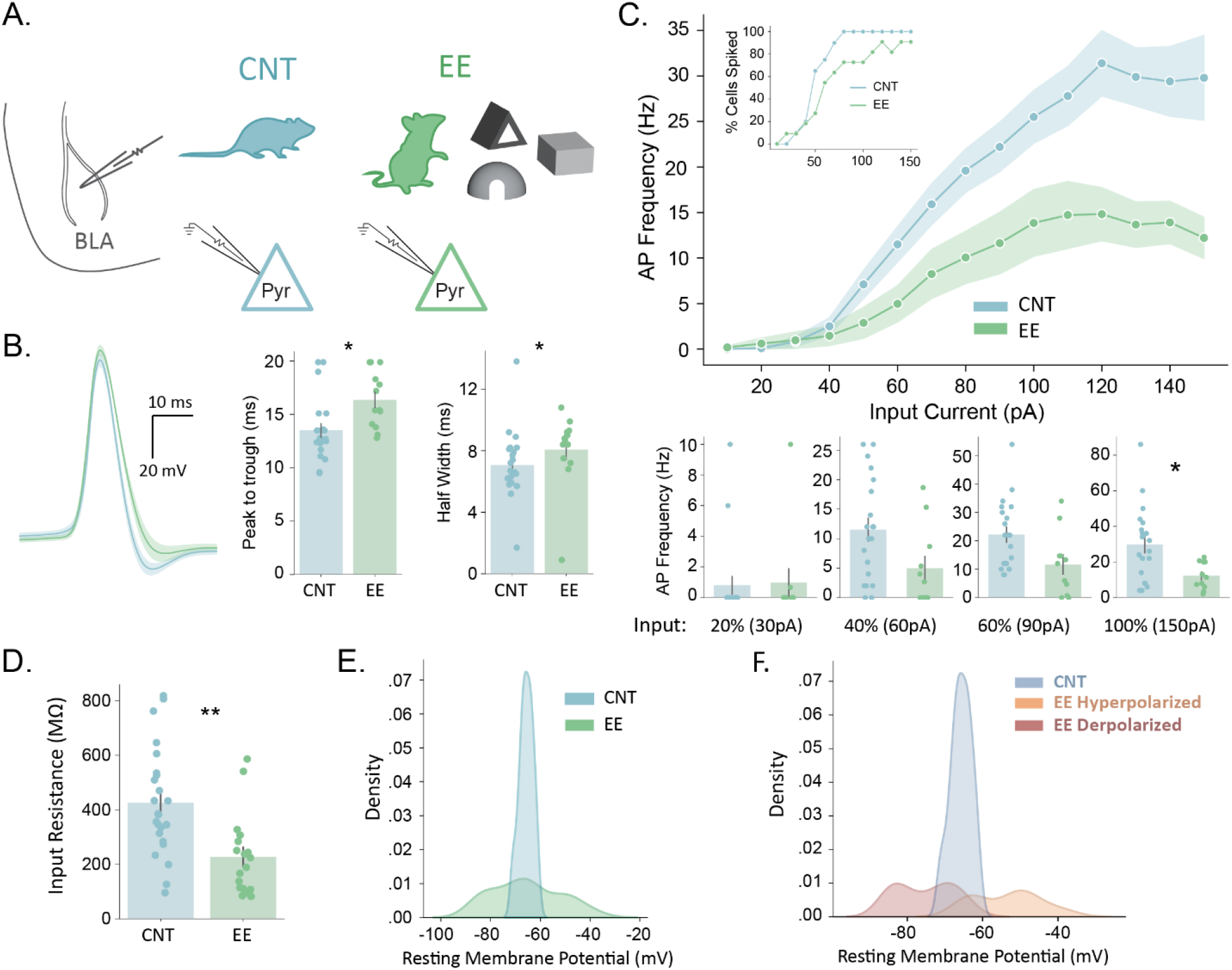
Enriched Environment exposure alters the electrophysiological properties of BLA principal neurons. **A)** Schematic diagram illustrating BLA principal cells recorded from control and EE mice. **B)** EE increases average action potential duration compared to controls. **C)** Input-output curve showing that EE reduces the maximum firing rate of principal cells. **D)** EE reduces average input resistance of BLA principal cells compared to controls. **E)** The variability of the resting membrane potential is higher in EE compared to control principal cells. **F)** Distribution plot illustrating putative depolarized and hyperpolarized clusters resulting from EE treatment, when split by the RMP mode of CNT cells. Assessed by permutation-based MANOVA and Mann-Whitney U

### Enriched environment alters the electrophysiological properties of BLA principal neurons in a projection-specific manner

Given the increased variability in the RMP of BLA principal neurons following EE, similar to what is observed following chronic stress, we investigated whether this maps onto distinct populations of projection neurons in the BLA. We used retrograde tracing to identify BLA-BnST and BLA-NAcc neurons and determine the impact of EE on their electrophysiological properties (Figure 5A). Unlike CUS, we found no difference in the shape of the AP or input-output features between BLA-BnST or BLA-NAcc neurons following EE (Figure 5B). Comparisons across electrophysiological properties between BLA-BnST and BLA-NAcc neurons from EE mice are shown in Supplemental Figure 6. Similar to CUS, we observed significant changes in the RMP of BLA-BnST and BLA-NAcc neurons following EE, although in the opposite direction with BLA-NAcc neurons being more depolarized (-60.13 ± 1.87 mV) compared to BLA-BnST neurons (-71.32 ± 1.34 mV) (Figure 5C). In the EE condition, the BLA-BnST neurons are not significantly different from the EE hyperpolarized population (- 75.30 ± 2.43 mV) and the BLA-NAcc neurons are not different from EE depolarized population (-53.56 ± 3.12 mV). Conversely, both BLA-NAcc and BLA-BnST neurons are significantly different from EE hyperpolarized and depolarized populations, respectively (Figure 5D), which is again opposite of the CUS condition. The linear classifier showed that the best predictor of BLA-BnST and BLA-NAcc populations after EE was, again, the separation of EE cells by the mode of the RMP in control cells (Figure 5E; Supplemental Table 2). Overall, our data suggest that EE induces changes in the electrophysiological properties of specific BLA principal neuron populations, promoting information flow from the BLA to NAcc whilst inhibiting information flow to the BnST.

**Figure 5:**
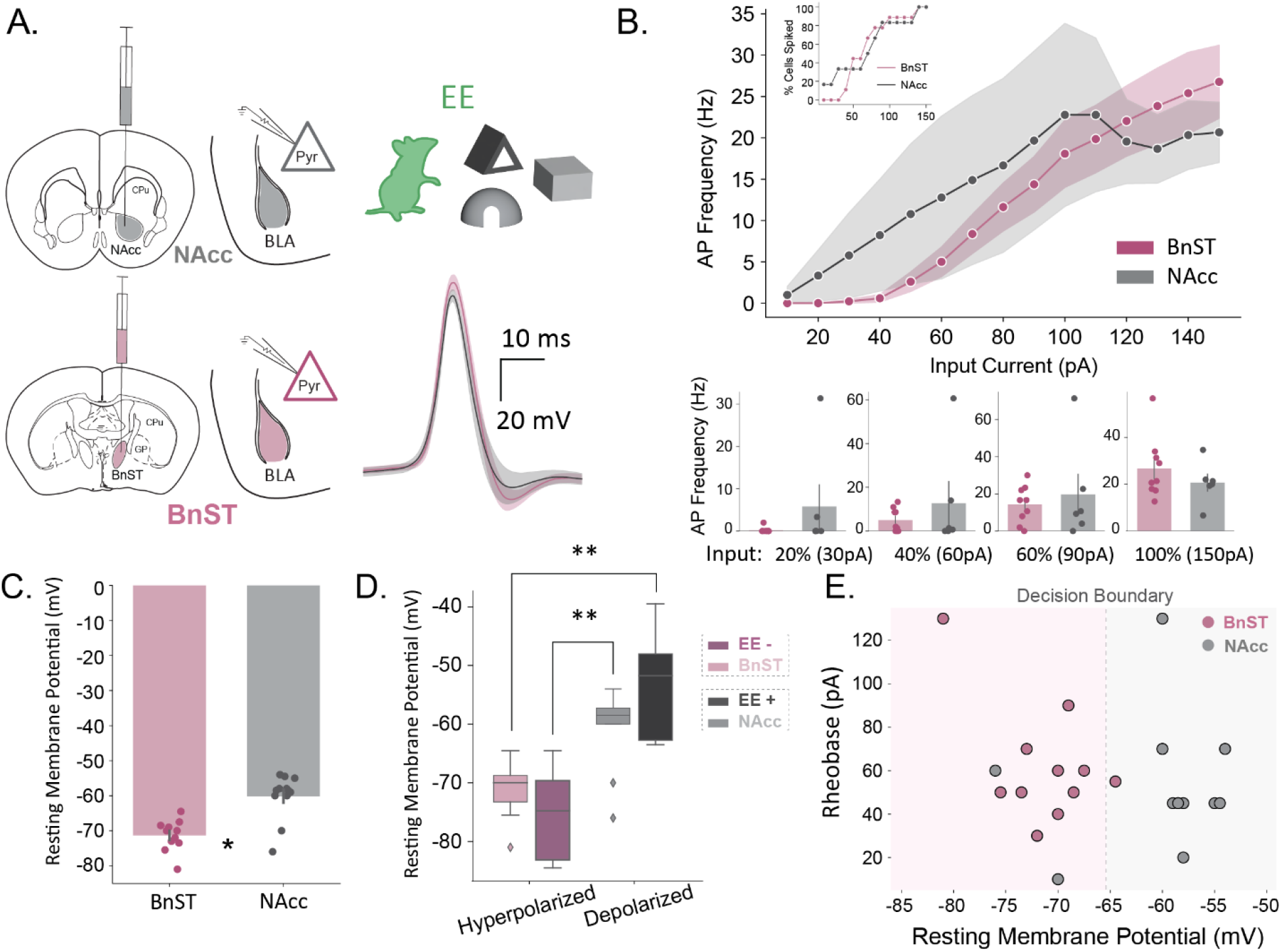
Projection-specific impact of enriched environment exposure on BLA principal neurons. **A)** Schematic diagram illustrating the targeting and recording of BLA principal cells projecting to either NAcc or BnST along with the average action potential waveform. **B)** Input-output curve (top), along with summary bar plots (bottom). **C)** The average resting membrane potential of BLA neurons projecting to either the BnST or NAcc. **D)** Comparison of resting membrane potential of NAcc and BnST cells with CUS hyperpolarized and depolarized clusters. **E)** Scatter plot of rheobase against resting membrane potential decision boundary of a gaussian naïve bayes classifier trained on EE depolarized and hyperpolarized clusters (Supplemental Table 2, row 0).

### Manipulating BLA information routing is sufficient to alter behavioral outcomes following enriched environment exposure

Our experiments show that EE may preferentially activate BLA-NAcc over BLA-BnST projections. As before we wanted to investigate if manipulating information routing through the BLA to specific downstream circuits is sufficient to alter behavioral outcomes associated with EE. We assessed CNT, EE, CNT BnST-NAcc+, and EE BnST+NAcc-mice across the OF, LD box, and TST behavioral paradigms (Figure 6A-B). We found that EE mice significantly increased the number of entries to the center of the OF and entries to the light zone of the LD box but had no effects on TST when compared to CNT mice (Figure 6C-E). EE mice also had higher total immobility in the OF than CNT mice, even though they covered the same total distance suggesting that they reach higher velocities when they are moving (Supplemental Figure 7A). These findings demonstrate that EE has a modest impact on behavioral states compared to controls which may be due to a floor effect since the behavior of control animals is not impaired. Interestingly, CNT BnST-NAcc+ mice exhibited a higher distance traveled, time in center, and entries in center of OF and reduced time immobile in the TST when compared to CNT mice (Figure 6C), suggesting that suppression of BLA-BnST projections and activation of BLA-NAcc projections reduces avoidance behaviors and stress-induced helplessness. EE BnST+NAcc-mice also exhibit robust differences across all behavioral paradigms when compared to EE mice, resulting in increased avoidance and stress-induced helplessness behaviors (Figure 6C-E). Indeed, the effect of activating BLA-BnST projections and suppressing BLA-NAcc projections was so robust that EE BnST+NAcc-almost fell entirely (81.8%) in a different cluster than all other groups (Figure 6F). These findings suggest that altering the information routing through BLA is sufficient to bidirectionally control behavioral outcomes following enriched environment exposure.

**Figure 6:**
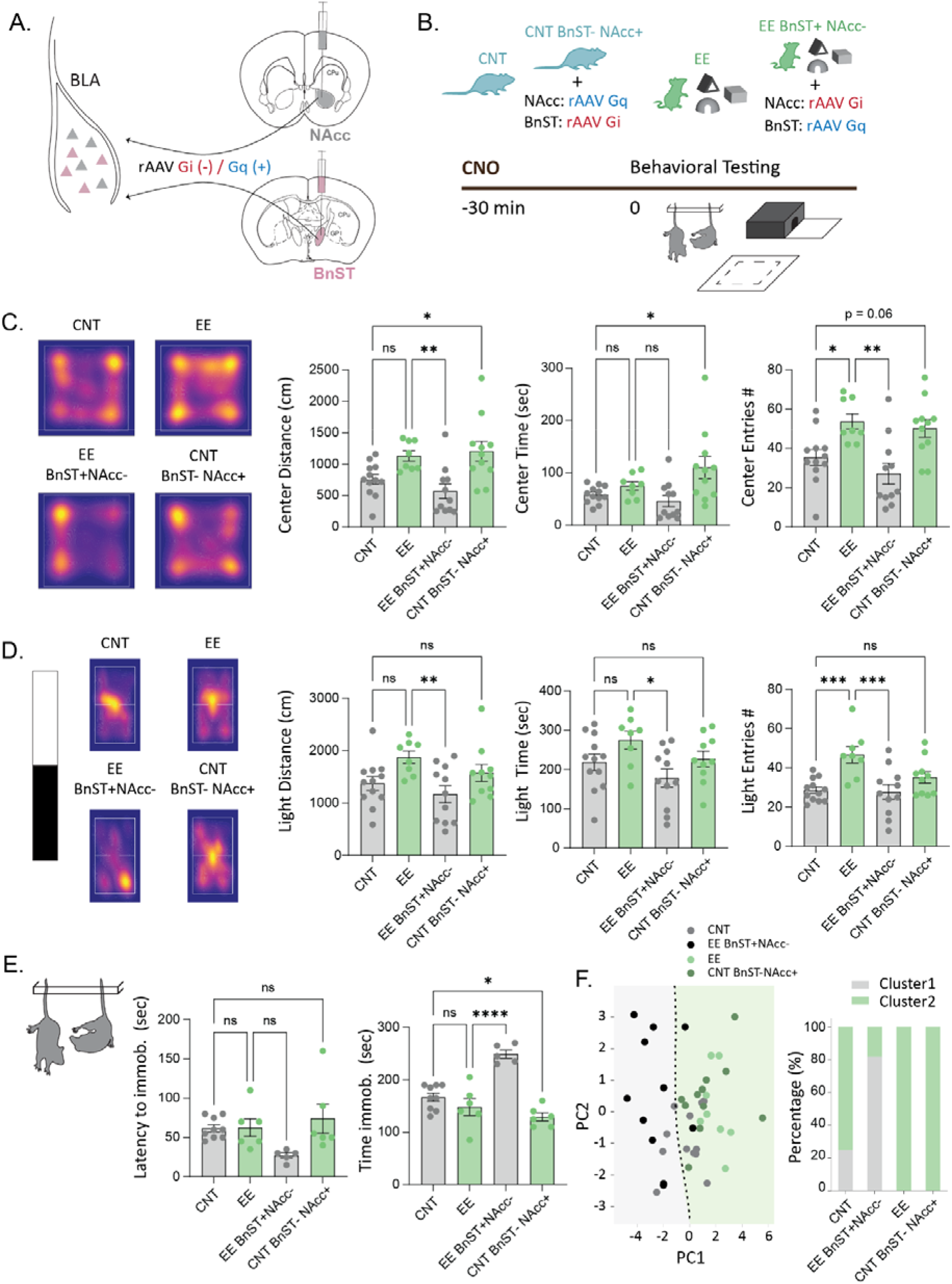
Manipulation of BLA-NAcc and BLA-BnST circuits alters behavioral outcomes following enriched environment. **A)** Schematic diagram illustrating targeting of BLA-BnST and BLA-NAcc pathway using retroAAV Gq/Gi DREADDs. **B)** Schematic diagram indicating the treatment groups (control: CNT, control with rAAV Gi in BnST and rAAV Gq in NAcc: CNT BnST-NAcc+, enriched-environment: EE, enriched-environment with rAAV Gq in BnST and rAAV Gi in NAcc: EE BnST+NAcc-), and that mice were injected with CNO (dose) 30 minutes prior to each behavioral test. **C)** Open field test representative heatmaps (left) and summary bar plots of distance, time, and entries in center zone across groups (right). **D)** Light Dark box test representative heatmaps (left) and summary bar plots of distance, time, and entries in light zone across groups (right). E) Tail suspension test summary bar plots quantifying latency to immobility and total time immobile. **F)** Clustering of the groups into two clusters using a GMM and the first five principal components. Decision boundary for the first 2 principal components (left) and percentage of each group per cluster (right).

### Previous experiences influence the reactivity and synchrony of subsets of BLA principal neurons

We used *in vivo* calcium imaging to quantify whether, and to what extent, context (CUS or EE paradigms) biases local BLA network activity. In vivo calcium imaging was recorded under baseline conditions where calcium (Ca) dynamics were assessed pre-and post-acute restraint stress (Figure 7A-B). Having experienced either CUS or EE for 4 weeks, recordings were again captured pre-and post-acute restraint stress, providing a means to compare the effect of previous experience on stress-evoked Ca responses in BLA. We recorded 1179 cells from CUS mice and 893 cells from EE mice and found no significant differences between the number of cells recorded at the collection time periods, pre-post restraint (Supplemental Figure 8A).

**Figure 7:**
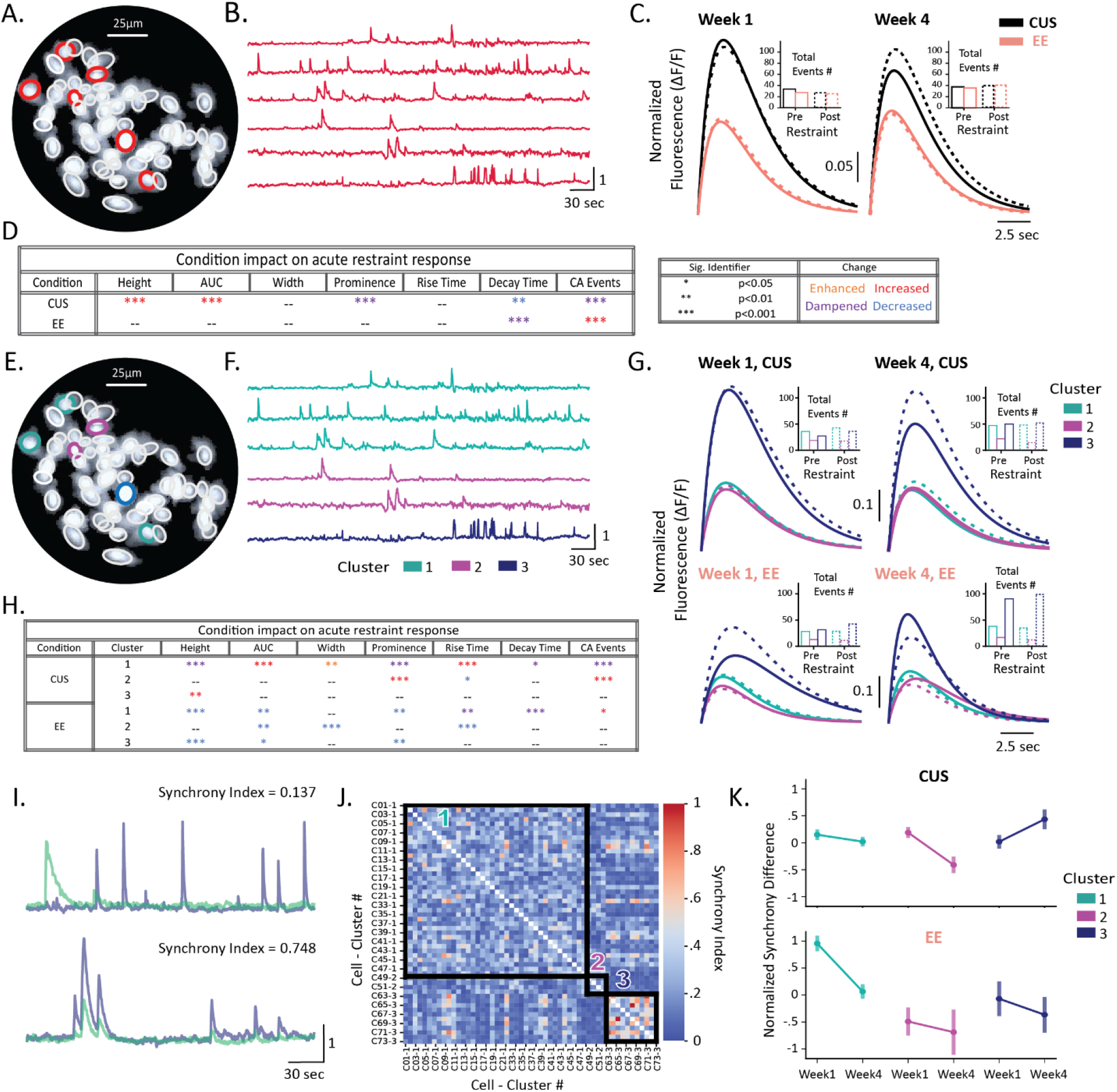
Context-specific changes in the reactivity and synchrony of principal neurons in the BLA. **A)** Representative image of a max projection FOV (field of view) showing GCaMP6f expression in mouse BLA. Scale bar: 25 μm. **B)** Ca^2+^ (Ca) traces of six representative BLA neurons that are shown in (**A**) encircled in red. **C)** Mean Ca waveforms from baseline (week 1; left) and post-context (week 4; right) recordings before (solid line) and after (dashed line) acute restraint. Inlay shows average Ca events for each condition; black: CUS, red: EE. **D)** Summary table detailing output of statistical tests (pooled independent t-tests on mean acute response differences) across each Ca-transient feature across CUS and EE conditions showing relatively little restraint-induced change for EE, as opposed to CUS. **E-F)** Same FOV image and representative traces shown in (**A** & **B**), recolored based on assigned cluster following unbiased gaussian mixture model (GMM). **G)** Mean Ca waveforms from baseline (week 1; left) and post-context (week 4; right) recordings before (solid line) and after (dashed line) acute restraint for the CUS (top) and EE (bottom) cohorts. Waveforms are colored with respect to cluster assignment detailed in **F)** with inlays showing average number of Ca events for each condition. **H)** Summary table detailing output of statistical tests (as shown in D), detailed by applied cluster **I)** Representative Ca traces for a set of cell pairs showing instances of low (top) and high (bottom) synchronization computed using instantaneous phase for each cell’s Ca signal (see Methods for more details). **J)** Synchronization matrix for representative recording (FOV and traces shown in **A/B** & **E/F**) shows pairwise synchrony indices for each cell recorded in single session with black squares outlining within (intra) cluster comparisons. Heat represents synchrony magnitude with warmer colors indicating high levels of synchronization. **K)** Stacked point-plots show restraint response difference in synchronization from baseline (week 1) and experience driven (week 4) by cluster; CUS-top, EE-bottom.

We then assessed how CUS and EE impacted Ca-transient features with respect to restraint. Our analysis revealed that context had a varied effect on restraint response for some of these features yet left the majority unaffected (Figure 7C and 7D). However, given that network states and concomitant behaviors might arise from independent BLA ensembles, it is possible that the lack of differences across context could derive from the assessment of aggregate ensemble activity (i.e. averaging across functionally-disparate cell populations). Consequently, the effects from specific sub-populations of principal cell clusters, each with different functional roles and potentially distinct Ca-transient profiles, could be masked by the mean differences observed across aggregate ensembles. To test this hypothesis, we subjected the data to an unbiased gaussian-mixture model (GMM) based on the Ca-transient features assessed in Figure 7C. A total number of 2018 (97.4% of total collected) cells were classified into three clusters (Figure 7E-F). The results further suggest 3 “types” of restraint-depended responses: cells that are (1) unchanged; (2) enhanced; or (3) reduced in both response magnitude and quantity. Within-cluster chi-square tests were performed and revealed that no significant differences in percentage of cells between baseline and EE/CUS, pre-post restraint (ps > 0.05, Supplemental Figure 8B). We observed that each cluster had a unique, context-dependent profile of feature changes, following acute restraint with CUS predominantly potentiating waveform attributes while EE attenuates them. Specifically, following CUS, cells in cluster 1 respond with slower and fewer events, while events in clusters 2 and 3 become larger and more frequent (Figure 7G and 7H). Conversely, EE reduces the magnitude of responses across all clusters, while increasing the frequency of events for cluster 1 responses (Figure 7G and 7H). These findings indicate that CUS and EE uniquely influence distinct BLA principal cell sub-populations, which may be responsible for conveying signals of positive and negative valence. However, the above analyses separated cells based on Ca-transient features and not on the functionality of each cluster.

To determine cluster distinctiveness, post-hoc, we calculated synchrony as a measure of functionality (Acebrón et al., 2005; Melloni et al., 2007) within (intra-) and between (inter-) clusters identified via the GMM. To this end, we quantified pair-wise synchronization (Patel, Tapan P. et al, 2015) from each recording (Figure 7I and 7J), positing that if the clusters are distinct, and functionally connected, the intra-cluster synchrony index would be higher than the inter-cluster index. This initial analysis revealed that synchronization within cluster was significantly higher than between cluster, regardless of context (Supplemental Figure 8C). We further found that CUS and EE impact synchronization in disparate ways. EE causes a homogenous response across all cell clusters by minimizing the effect of acute restraint on synchrony—doing so significantly for cluster 1. Conversely, CUS causes a reduction in synchrony following acute restraint for clusters 1 and 2, meanwhile magnifying synchrony following acute restraint for cluster 3 (Figure 7K). These results suggest that the GMM identified ensembles of functional clusters which are impacted distinctly based on animal experience; a finding that is in accordance with our behavioral and physiology results.

## Discussion

These data demonstrate a novel mechanism through which previous experiences influence valence processing by altering the electrophysiological properties of projection-specific neurons and, thereby, the excitability of specific subsets of principal neurons in the BLA, biasing information routing through the BLA, and directing information flow to specific downstream circuits. We demonstrate that chronic stress facilitates the activity of BnST-projecting principal neurons in the BLA while suppressing the activity of NAcc-projecting BLA neurons. Mimicking this biased information routing using simultaneous chemogenetic activation of BLA-BnST neurons and suppression of BLA-NAcc neurons, we are able to phenocopy the behavioral deficits associated with chronic stress in naïve mice. Conversely, we are able to mitigate the behavioral deficits following chronic stress by simultaneously activating BLA-NAcc neurons and suppressing BLA-BnST neurons. We also demonstrate that positive experiences bias information routing through the BLA, facilitating information flow to the NAcc and impeding information transfer to the BnST. Chemogenetic manipulations that mimic this pattern of information flow (activating BLA-NAcc and suppressing BLA-BnST) is sufficient reduce avoidance and stress-induced helplessness behaviors. Conversely, chemogenetic activation of BLA-BnST and suppression of BLA-NAcc in EE mice can enhance avoidance and stress-induced helplessness behaviors. These data suggest that biased information routing through the BLA due to altered electrophysiological properties of projection-specific BLA principal neurons can bidirectionally control valence processing and thus the outcomes on avoidance and stress-induced helplessness behaviors. While the electrophysiological and chemogenetic experiments tell us how modulation of projection cells influences behavioral outcomes, they are limited in their ability to infer how network activity in BLA is impacted by experience.

To examine how BLA networks are impacted by previous experience, we employed in vivo calcium (Ca) imaging in mice in response to acute stress following CUS or EE. Notably, our in vivo imaging analyses were not solely based on Ca-transient features but also considered the timing of these features. By examining intra-and inter-cluster synchrony, we confirmed the distinctiveness of each cluster, and these results indicate that CUS and EE differentially impact both the transient properties and functional connections of distinct BLA principal cell sub-populations, potentially contributing to the different influences of these experiences on affective states. These findings have several implications for understanding the neural mechanisms underlying stress and resilience. The observed differences in BLA principal cell responses to CUS and EE shed light on how different environmental contexts may shape emotional processing and responses to stress. The distinct clusters identified by the GMM suggest that specific functional ensembles of principal cells may be responsible for conveying positive and negative affective signals within BLA. Additionally, investigating the downstream effects of BLA signaling on other brain regions involved in emotion regulation could provide a more comprehensive understanding of the broader neural circuitry involved in stress, resilience, and valence processing.

These data show that previous experiences shape the activity of principal neurons in the BLA in a projection-specific manner, through a mechanism involving altered electrophysiological properties of specific subsets of BLA principal neurons. This plasticity mechanism likely alters the participating neurons during ensemble recruitment, which relies on the underlying excitability of principal neurons in the BLA (Josselyn and Frankland, 2018; Lau et al., 2020; Lisman et al., 2018; Rashid et al., 2016; Yiu et al., 2014). Thus, previous experiences may influence future reactivity to emotionally salient stimuli, such as fear, due to altered intrinsic excitability of projection-specific BLA principal neurons. This may represent a mechanism whereby salient experiences exert long-term impacts on mood. Future studies are required to determine whether a similar pathological mechanism may play a role in exacerbated fear reactivity, fear generalization, and potentially to the mechanisms mediating PTSD.

The mechanisms mediating the changes in intrinsic excitability of projection-specific BLA principal neurons following either chronic stress or EE remain unclear, although it appears to involve changes in ion channels responsible for setting the resting membrane potential of these neurons. This is a critical remaining question regarding the cellular and molecular basis of the electrophysiological changes underlying the biased information routing following either positive or negative experiences. Previous studies have demonstrated that chronic corticosterone administration increases negative valence processing and impairs positive valence processing (Dieterich et al., 2019), implicating corticosterone and glucocorticoid signaling in this process. Previous studies have also demonstrated that acute stress alters information processing in the PFC biasing information flow from the BLA over other inputs through an mGlu5-dependent plasticity mechanism (Joffe et al., 2022). Albeit mechanistically different from the changes in intrinsic excitability mediating chronic stress-induced biased information routing in the BLA, these studies demonstrate that stress can influence information flow through circuits involved in emotional processing. Future studies are required to understand the precise mechanisms mediating the impact of chronic stress or enriched environment on the electrophysiological properties of principal neurons in the BLA and information routing to downstream circuits. Despite this gap in our current knowledge, the data presented in this manuscript suggests that this mechanism may contribute to long-term changes in processing emotionally salient information and may contribute to the allostatic load leading to maladaptive behaviors (McEwen, 1998).

It is important to note that both the BnST (Lebow and Chen, 2016) and NAc (Ray et al., 2022; Zhou et al., 2022) have been shown to contribute to both positive and negative valence processing. In fact, BLA projections to the NAc has been shown to promote positive reinforcement via disinhibiting dopamine neurons in the ventral tegmental area (VTA) (Zhou et al., 2022), in contrast to our observations in the current study where activating BLA-NAcc reduces avoidance behaviors and stress-induced helplessness. It is likely that even within the BLA-NAcc projections there are valance-specific neurons and that our current approach is not sufficient to resolve this nuance within the circuit. Further, our chemogenetic manipulations involve the simultaneous manipulation of both BLA-NAc and BLA-BnST neurons making it difficult to resolve the impact of a singular circuit on the behavioral outcomes, although the simultaneous recruitment and/or suppression of multiple circuits in parallel is likely more physiologically relevant. Finally, it is possible that the role of these specific circuits is altered by previous experiences, such as chronic stress fundamentally altering how they participate in behavioral outcomes.

Extensive studies have investigated the participation of specific ensembles and specific circuits in valence processing. To our knowledge, this is the first study to examine how these circuits are fundamentally altered under pathological conditions, such as chronic stress, to bias circuit engagement and influence behavioral outcomes. Further, we demonstrate an opposing impact of enriched environment on information flow through the BLA and valence processing. These studies contribute to our knowledge of the cellular and molecular mechanisms through which previous experiences can exert long-term impacts on emotional processing.

## Methods

### Animals

Adult (10-12 weeks old) male C57BL/6J (Jackson Labs; stock # 000664) were purchased directly from The Jackson Laboratory. Mice were housed at Tufts University School of Medicine’s Division of Laboratory Animal Medicine facility in clear plastic cages (4 mice per cage), in a temperature and humidity-controlled facility, on a 12 hour light/dark cycle (lights on at 0700 h), with ad libitum access to food and water for at least a week prior to experimentation. All procedures were approved by the Tufts University Institutional Animal Care and Use Committee.

### Chronic unpredictable stress paradigm

Adult C57BL/6J mice underwent a three-week CUS protocol as previously described (Antonoudiou et al., 2022; Walton et al., 2023). Briefly, mice were subjected to four consecutive days of alternating stressors only during the dark period, including cage tilt, restricted cage, food and water restriction, soiled cage, and unstable cage, followed by an acute stressor on the fifth day (restraint stress, 2-hours of cold stress exposure or tail suspension) each week for 3 consecutive weeks. Mice were minimally handled in parallel to serve as controls.

### Enriched environment paradigm

Adult C57BL/6J mice were housed in a homecage with enriched environment compared to the standard housing conditions. Mice were housed in a large mouse cage (10 1⁄2” x 19” x 8”) containing toys and habitat accessories (Kaytee cage accessories), a dam house, and an increased number of mice per cage (10 mice per cage) for 3 consecutive weeks. Mice housed in standard housing conditions served as controls.

### Retrograde labeling

Mice were anesthetized with an intraperitoneal injection of a ketamine/xylazine cocktail (100 mg/kg ketamine and 10 mg/kg xylazine) and administered sustained release buprenorphine (0.5mg/kg, subcutaneous injection) prior to surgery. Mice were stereotaxically injected with 250nl of AAVrg-hSyn1-GCaMP6f-P2A-nls-dTomato (Addgene #51085-AAVrg) into the NAcc (AP: +1.70; ML: +/- 0.75; DV: -4.0) or BnST (AP: +0.02; ML: +/- 0.50; DV: -4.0) using a 33-gauge Hamilton syringe at an infusion rate of 100nL/minute. Mice were allowed to recover for 3 weeks prior to experimentation to allow for optimal viral expression.

### Electrophysiology

Mice were anesthetized with isoflurane, decapitated, and the brain was rapidly removed and placed immediately in ice-cold, oxygenated normal artificial cerebrospinal fluid [nACSF; containing (in mM) 126 NaCl, 26 NaHCO_3_, 1.25 NaH_2_PO_4_, 2.5 KCl, 2 CaCl_2_, 2 MgCl_2_, and 10 dextrose (300–310 mOsm)] containing 3 mM kynurenic acid and bubbled with 95% O_2_-5% CO_2_. Coronal hippocampal slices (350 μm thick) were prepared using a Leica VT1000S vibratome and allowed to recover for at least 1 hour prior to recording. Electrophysiological recordings were performed in a recording chamber maintained at 33°C (in-line heater; Warner Instruments) and perfused at a high flow rate (∼4 ml/min) throughout the experiment. Intrinsic electrophysiological properties were measured in visually-identified BLA principal neurons and in projection-specific neurons identified by expression of a retrograde fluorescent tag (GFP or tdTomato). Series resistance and whole cell capacitance were continually monitored and compensated throughout the course of the experiment. Recordings were eliminated from data analysis if series resistance increased by >20%. For all electrophysiology experiments, data acquisition was carried out using an Axopatch 200B (Axon Instruments) and PowerLab hardware and software (ADInstruments).

Data analysis was performed using Python analysis scripts developed in-house. Action potentials (APs) were detected using SciPy’s *find_peaks* function (prominence=50 mV, wlen=100 ms, distance=1 ms). For input-output (IO) curve analysis, only cells that had more than 10 Hz maximum firing rate (5 APs in 0.5 seconds), were included. The IO slope was calculated using a first-degree least squares polynomial fit (NumPy’s *polyfit*, n=1) between rheobase to 80% of max firing rate. For the IO slope, cells were included only if they fired for at least 3 steps from rheobase to 80% of max APs (at least 3 points for line fit). For waveform analysis, only a maximum of 3 APs were collected from up to the first 3 current steps above rheobase. This was done, to ensure that the afterhyperpolarization (AHP) was maximally preserved (Supplemental Figure 2A). For all analyses, cells were excluded based on their maximum firing rate using Tukey’s outlier detection method (k = 3). To calculate waveform properties, APs were interpolated (SciPy’s *interp1d*, interpolation factor=10, type=cubic). The AP threshold was detected as 0.15 mV/ms, the AP amplitude was defined as the difference from the AP peak – threshold. The AHP amplitude was defined as the absolute difference between the AHP peak – AP threshold. AP peak-to-trough was calculated as the time between AHP peak to AP peak. The rise time was calculated as the time between AP peak to AP threshold.

To determine the best predictor(s) for identifying BLA projections to BnST and NAcc after CUS and EE (testing data), we trained a gaussian naïve bayes (GNB) linear classifier using a combination of features from the CUS and EE non-specific projections (training data) as outlined in Supplemental Tables 1 and 2. Features were always z-normalized at the beginning of the pipeline. For each set of features, (each row in the Supplemental Tables 1 and 2) cells that were containing empty features were excluded. Features were split into two groups based on either the mode of the selected features (most frequent observation) or by feeding the features into a Gaussian Mixture Model (GMM). Features were fed into the GMM as is, or by performing dimensionality reduction using principal component analysis (PCA; sklearn’s *decomposition.PCA*). Before performing PCA, highly correlated features were removed (Pearson correlation coefficient > .95) and the number of principal components (PCs) for feeding into the GMM was selected based on cumulative explained variance threshold (Supplemental Tables 1 and 2). For each set of features the *covariance matrix* that resulted in the best silhouette score was used for clustering the data using the GMM. Further, for each set of features a GNB model (sklearn’s *naive_bayes*) was fined tuned by adjusting the var_smoothing parameter and testing its performance using gridsearch (sklearn’s *GridSearchCV*: scoring=balanced_accuracy) and cross-validation (sklearn’s *StratifiedKFold*: nsplits=3). Each GNB model was then trained on 90% of the training data and generated predictions for group clusters that were assessed using Balanced Accuracy and F1 score using the testing data. For the purposes of visualization of the scatter decision plots (Figure 2F, 5E), missing rheobase values were imputed using k-Nearest Neighbors (sklearn’s *KNNImputer*).

### In vivo calcium imaging

Mice were anesthetized with an intraperitoneal injection of a ketamine/xylazine cocktail (100 mg/kg ketamine and 10 mg/kg xylazine) and administered sustained release buprenorphine (0.5mg/kg, subcutaneous injection) prior to surgery. Mice were stereotaxically injected with 500nl of AAV.CamKII.GCaMP6f.WPRE.SV40 (Addgene # 100834) into the BLA (AP: -1.50; ML: +/- 3.0; DV: -4.5) using a 33-gauge Hamilton syringe at an infusion rate of 100nL/minute. After GCaMP infusion, an Inscopix lens affixed to a baseplate (Inscopix, 1050-004413) was placed in the BLA. Mice were allowed to recover for 6 weeks prior to experimentation to allow for optimal viral expression and *in vivo* imaging.

On the day prior to imaging experiments, animals were moved from the animal housing facility to the room where experimental procedures took place, limiting undue stress prior to baseline recordings. On the first day of recordings, each animal was removed from their cage, had a miniature integrated microscope system (nVoke HD 2.0; Inscopix) attached to their baseplate, and was given 10 min to habituate to the testing cage (a cage identical to homecage, containing fresh bedding/nesting). Images were acquired using data acquisition software (ver. 2.0.0; Inscopix) at 20 frames per sec, 20% of LED power, and a gain kept between (2 and 5, varying based on clarity of field of view; FOV). Images were acquired for 10 min pre-and post-acute restraint stress (30min in modified, clear polypropylene Falcon tubes (Melón et al., 2017)), after which the microscope was demounted from the baseplate, and animal was returned to their home-cage. Animals were subjected to 3 weeks of either chronic unpredictable stress (CUS) or enriched environment (EE), at which point, the previously described protocol was repeated. In total, each animal provided four recordings: pre-and post-acute restraint in weeks 1 and 4.

Acquired imaging data were down-sampled (1/2 spatial binning), preprocessed, motion corrected, and then calcium transients of individual neurons were extracted with a constrained non-negative matrix factorization for microendoscopic data (CNMF-E) using Inscopix Data Processing Software (ver. 1.9.1; Inscopix). All extracted traces were manually checked resulting in traces from multiple cells or non-cellular signals being excluded. The fluorescent trace data was then subjected to a series of custom-built scripts (implementing SciPy’s (Virtanen et al., 2020) peak-finding algorithms in Python) which automated the detection of calcium (CA) transient events, generating a structure which detailed the peak amplitude (height), area under the curve (AUC), width, peak-to-peak prominence (prominence), rise time, and decay time for each traces’ events within a given recording.

A clustering analysis was used to identify groups of neurons that were impacted similarly across experimental conditions. In brief, the detailed structure was normalized (CA features from all collected traces were minmax scaled) and the number of clusters were unbiasedly determined based on the best fit using a Gaussian Mixture Model (GMM; calculating the Bayesian information criterion) as done previously in (Athey and Vogelstein, 2019; Stone et al., 2022). Classifications were considered valid only if they fell within the 90% confidence interval from the centroid of each respective cluster, after which we constrained our analyses to neurons within the same cluster.

Network synchronization was evaluated using custom-built scripts in Python, as employed in recent *in vivo* imaging studies (Liu and Baraban, 2019; Patel et al., 2015). In brief, a pair-wise phase synchronization index was computed for every combination of cells in a given recording session, denoting likelihood of neuronal assemblies being functional units (i.e. ensembles) via a value range from 0 (non-coordinated activity) to 1 (completely synchronized activity). These values were then normalized (by recording) to account for cases in which there were disproportionate ratios of within-cluster (intra) to between-cluster (inter) comparisons. This provides a means to accurately compare network synchronization between conditions, clusters, and cohorts. The resulting normalized values represent the change-fold difference between intra-and inter-cluster synchrony, with values greater than 1 indicating elevated cluster coherence. Note: this synchronization analysis assumes that the state of the BLA did not change over the course of the 10-minute imaging session.

### Chemogenetics

Mice were anesthetized with an intraperitoneal injection of a ketamine/xylazine cocktail (100 mg/kg ketamine and 10 mg/kg xylazine) and administered sustained release buprenorphine (0.5mg/kg, subcutaneous injection) prior to surgery. Information routing through the BLA was manipulated using chemogenetics to mimic or prevent the effects of CUS or EE:

#### DREADD CUS

To mimic CUS, mice were stereotaxically injected with AAVrg-EF1a-Cre (Addgene #55636) in the NAc (AP: +1.70; ML: +/- 0.75; DV: -4.0), AAVrg-hSyn-fDIO-hM3D(Gq)-mCherry-WPREpA (Addgene #154868) in the BnST (AP: +0.02; ML: +/- 0.50; DV: -4.0), and AAV1-EF1a-Flpo (Addgene #55637) and AAV8-hSyn-DIO-HA-hM4D(Gi)-IRES-mCitrine (Addgene #50455) in the BLA (AP: -1.50; ML: +/- 3.0; DV: -4.5) using a 33-gauge Hamilton syringe at an infusion rate of 100nL/minute. To prevent the effects of CUS, mice were injected with AAVrg-hSyn-fDIO-hM3D(Gq)-mCherry-WPREpA in the NAcc, AAVrg-EF1a-Cre in the BnST, and AAV1-EF1a-Flpo and AAV8-hSyn-DIO-HA-hM4D(Gi)-IRES-mCitrine in the BLA and subsequently subjected to the 3-week CUS paradigm.

#### DREADD EE

To mimic the effects of EE, mice were stereotaxically injected with AAVrg-hSyn-fDIO-hM3D(Gq)-mCherry-WPREpA in the NAcc, AAVrg-EF1a-Cre in the BnST, and AAV1-EF1a-Flpo and AAV8-hSyn-DIO-HA-hM4D(Gi)-IRES-mCitrine in the BLA. To prevent the effects of EE, mice were stereotaxically injected with AAVrg-EF1a-Cre in the NAcc, AAVrg-hSyn-fDIO-hM3D(Gq)-mCherry-WPREpA in the BnST, and AAV1-EF1a-Flpo and AAV8-hSyn-DIO-HA-hM4D(Gi)-IRES-mCitrine in the BLA and then subjected to the 3-week EE protocol.

Following 3 weeks to allow for optimal DREADD expression and the conclusion of either the CUS or EE protocol, avoidance behaviors and stress-induced helplessness was assessed in these animals 30-min following an injection with 3mg/kg clozapine-n-oxide (CNO).

### Behavioral approaches

Behavioral assessments were conducted as previously described by our laboratory (Antonoudiou et al., 2022; Melón et al., 2017).

#### Open Field (OF)

Mice were placed in the center of the open field (40 cm x 40 cm) apparatus and the number of entries, distance traveled, and total time spent in the center of the open field was measured over a 10-min period using automated Motor Monitor software (Hamilton-Kinder).

#### Light/Dark Box (LD)

Mice were placed into the dark chamber of the two-chamber apparatus which is enclosed by a 22 x 43 cm photobeam frame with equally spaced photocells. The number of entries, distance traveled, and the amount of time spent in the light chamber was measured using automated Motor Monitor software (Hamilton-Kinder) over a 10-min period.

#### Tail Suspension (TST)

Mice were suspended by their tail on a bar 36 inches above the ground for a 6-min period and the latency to immobility and the total time immobile was measured.

To cluster behavioral data, all 12 behavioral features (5 OF, 5 LD, 2 TST) were used. Mice that were missing more than half of the behavioral features (2 out 3 behavioral tests) were excluded from further analysis. If any remaining mice had missing features, those features were imputed based on the median value of that feature according to its treatment group. The features were then z-normalized, and dimensionality reduction was performed using a PCA. The number of PCs that accounted for a cumulative explained variance ratio of up to 0.9 of principal components were selected for clustering using a GMM. The number of clusters (range = 2 - 4 clusters) and covariance matrix of the GMM was selected based on the best silhouette score.

### Statistical methods

Statistical analyses were performed using GraphPad Prism software, and Python (Python Software Foundation). For electrophysiological parameters between two groups Mann-Whitney U tests were run with Benjamini/Hochberg procedure to control for False Discovery Rate (statsmodels). To compare between clusters a permutation-based multivariate analysis of variance (MANOVA) was performed in python. To compare the resting membrane potential between the groups Kruskal-Wallis with Dunn multiple comparisons was run (PRISM). For comparison of behavioral groups, one-way ANOVAs with Sidak’s multiple comparisons were performed. For in-vivo calcium data, unless otherwise noted, the statistical package Pingouin was used. Chi-square tests were implemented to test condition*week and condition*cluster cell quantities. T-tests on pooled means and standard deviations were used to compare response differences due to acute restraint for condition*week and condition*week*cluster. Values are reported as the mean ± SEM. Error bars and shaded regions represent SEM. A p-value <0.05 was considered statistically significant.

