## Supplemental Material for "Experience-dependent information routing through the basolateral amygdala"

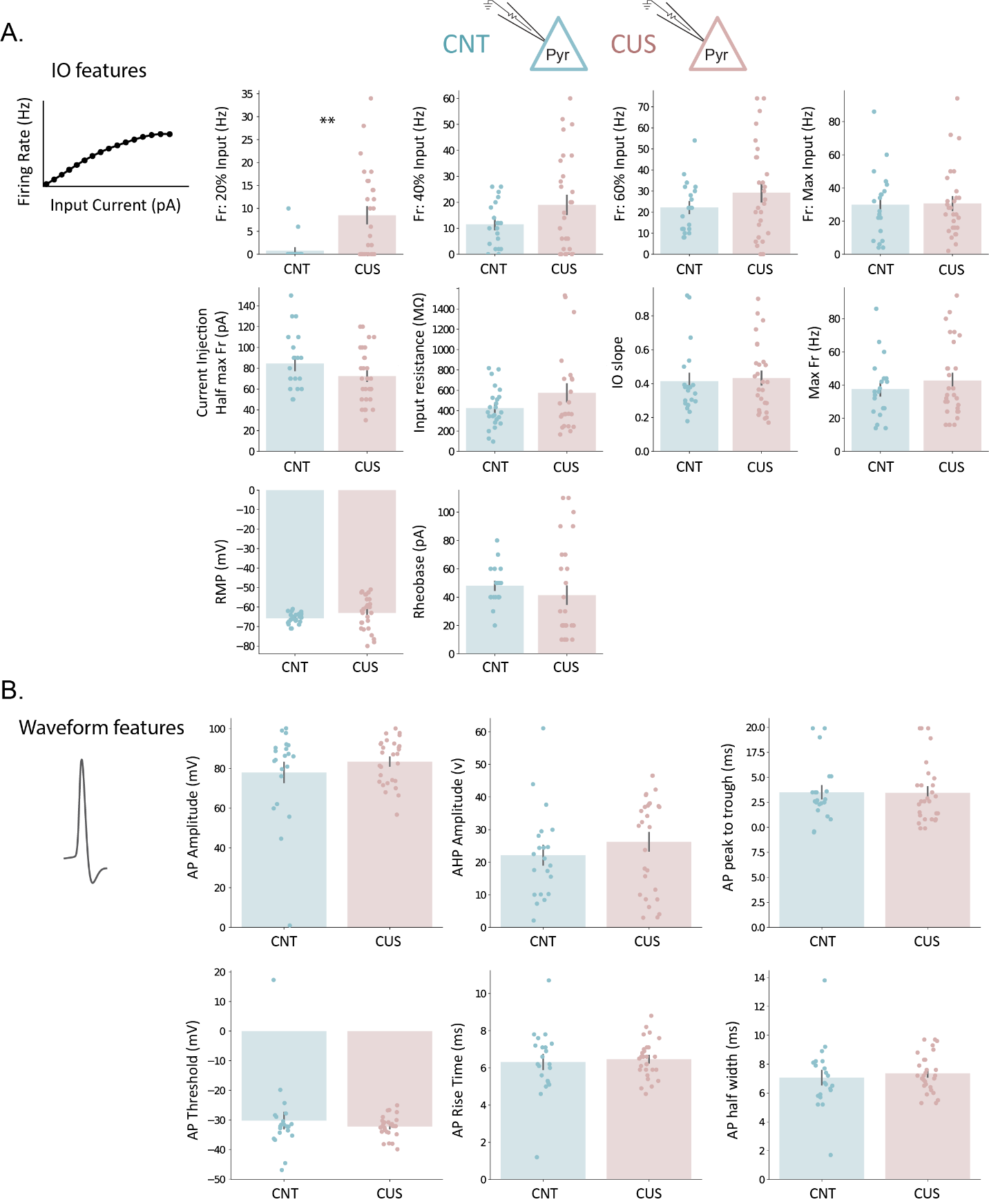

**Supplemental Figure 1: Effect of CUS on BLA principal cell properties.** **A)** Bar plots with individual points for input-output (IO) properties/features. **B)** Bar plots with individual points for action potential waveform properties/features.

| **Description** | **Figure** | **Stat test** | **Stats report** |
| --- | --- | --- | --- |
| Fr. 20% input | Figure 1B, Supp. Figure 1A | Mann–Whitney U & BH correction | U=129.5, p=0.004, Ncontrol=20, Ncus=28, |
| Fr. 40% input | Figure 1B, Supp. Figure 1A | Mann–Whitney U & BH correction | U=244.0, p=0.709, Ncontrol=20, Ncus=28, |
| Fr. 60% input | Figure 1B, Supp. Figure 1A | Mann–Whitney U & BH correction | U=247.0, p=0.709, Ncontrol=20, Ncus=28, |
| Fr. Max input | Figure 1B, Supp. Figure 1A | Mann–Whitney U & BH correction | U=282.0, p=0.975, Ncontrol=20, Ncus=28, |
| Current Inj. at Half Max Fr | Supp. Figure 1A | Mann–Whitney U & BH correction | U=343.0, p=0.471, Ncontrol=20, Ncus=28, |
| Input Resistance | Supp. Figure 1A | Mann–Whitney U & BH correction | U=237.5, p=0.709, Ncontrol=24, Ncus=23, |
| IO slope | Supp. Figure 1A | Mann–Whitney U & BH correction | U=257.0, p=0.876, Ncontrol=20, Ncus=27, |
| Max Fr. Rate | Supp. Figure 1A | Mann–Whitney U & BH correction | U=255.5, p=0.769, Ncontrol=20, Ncus=28, |
| Resting Membrane Potential | Supp. Figure 1A | Mann–Whitney U & BH correction | U=242.0, p=0.286, Ncontrol=24, Ncus=28, |
| Rheobase | Supp. Figure 1A | Mann–Whitney U & BH correction | U=364.5, p=0.286, Ncontrol=20, Ncus=28, |
| AP amplitude | Supp. Figure 1B | Mann–Whitney U & BH correction | U=271.0, p=0.919, Ncontrol=21, Ncus=28, |
| AHP amplitude | Supp. Figure 1B | Mann–Whitney U & BH correction | U=236.0, p=0.919, Ncontrol=21, Ncus=28, |
| AP peak to trough | Supp. Figure 1B | Mann–Whitney U & BH correction | U=307.0, p=0.919, Ncontrol=21, Ncus=28, |
| AP threshold | Supp. Figure 1B | Mann–Whitney U & BH correction | U=300.0, p=0.919, Ncontrol=21, Ncus=28, |
| AP rise time | Supp. Figure 1B | Mann–Whitney U & BH correction | U=288.5, p=0.919, Ncontrol=21, Ncus=28, |
| AP half width | Supp. Figure 1B | Mann–Whitney U & BH correction | U=250.5, p=0.919, Ncontrol=21, Ncus=28, |
| input-output properties, Control-clusters | Figure 1D | Permutation-based MANOVA | pseudo-F(1, 18) = 0.345, p < 0.888 |
| wave properties, Control-clusters | Figure 1D | Permutation-based MANOVA | pseudo-F(1, 19) = 1.760, p < 0.165 |
| input-output properties, CUS-clusters | Figure 1D | Permutation-based MANOVA | pseudo-F(1, 21) = 3.769, p < 0.041 |
| wave properties, CUS-clusters | Figure 1D | Permutation-based MANOVA | pseudo-F(1, 26) = 3.859, p < 0.014 |

**Stats Table 1.** Highlighted rows: p < 0.05

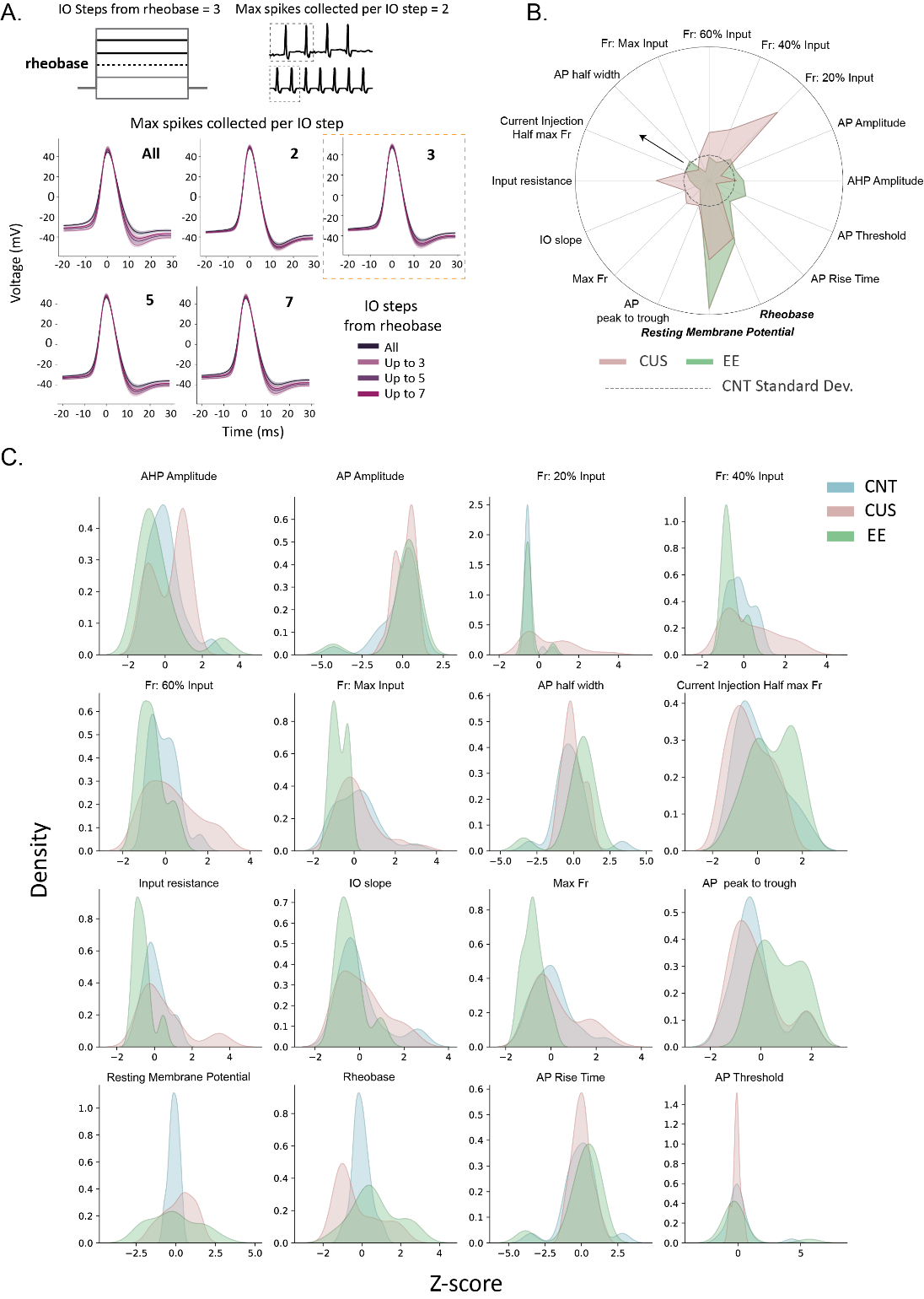

**Supplemental Figure 2: Examination of electrophysiological properties.**  **A)** Selection of action potentials (APs) from the input-output protocol that best preserves action potential amplitude and afterhyperpolarization. Orange square denotes the parameters used for AP collection. **B)** Radar plot showing the standard deviation of all extracted properties in CUS and EE animals normalized to CNT standard deviation represented as a grey dotted circle. Anything outside the dotted circle towards the direction of the arrow illustrates higher standard deviation than controls. Resting membrane potential and rheobase standard deviation consistently increase from CNT in both CUS and EE conditions. **C)** Distribution plot of all extracted properties in control, CUS and EE animals.

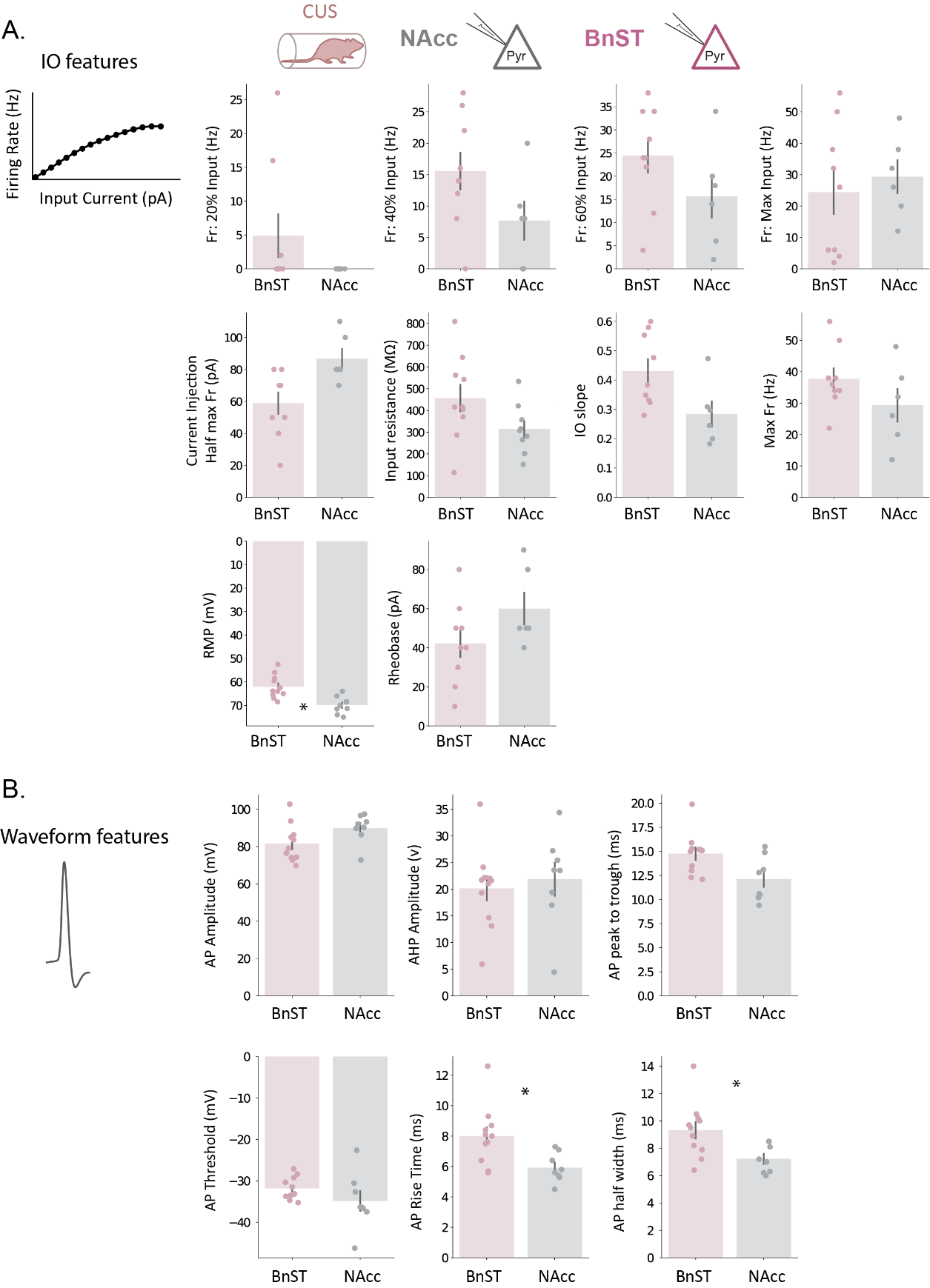

**Supplemental Figure 3:** **Projection-specific effects of CUS on BLA principal cell properties.** **A)** Bar plots with individual points for input-output (IO) properties/features in BLA-BnST and BLA-NAcc populations. **B)** Bar plots with individual points for action potential waveform properties/features in BLA-BnST and BLA-NAcc populations.

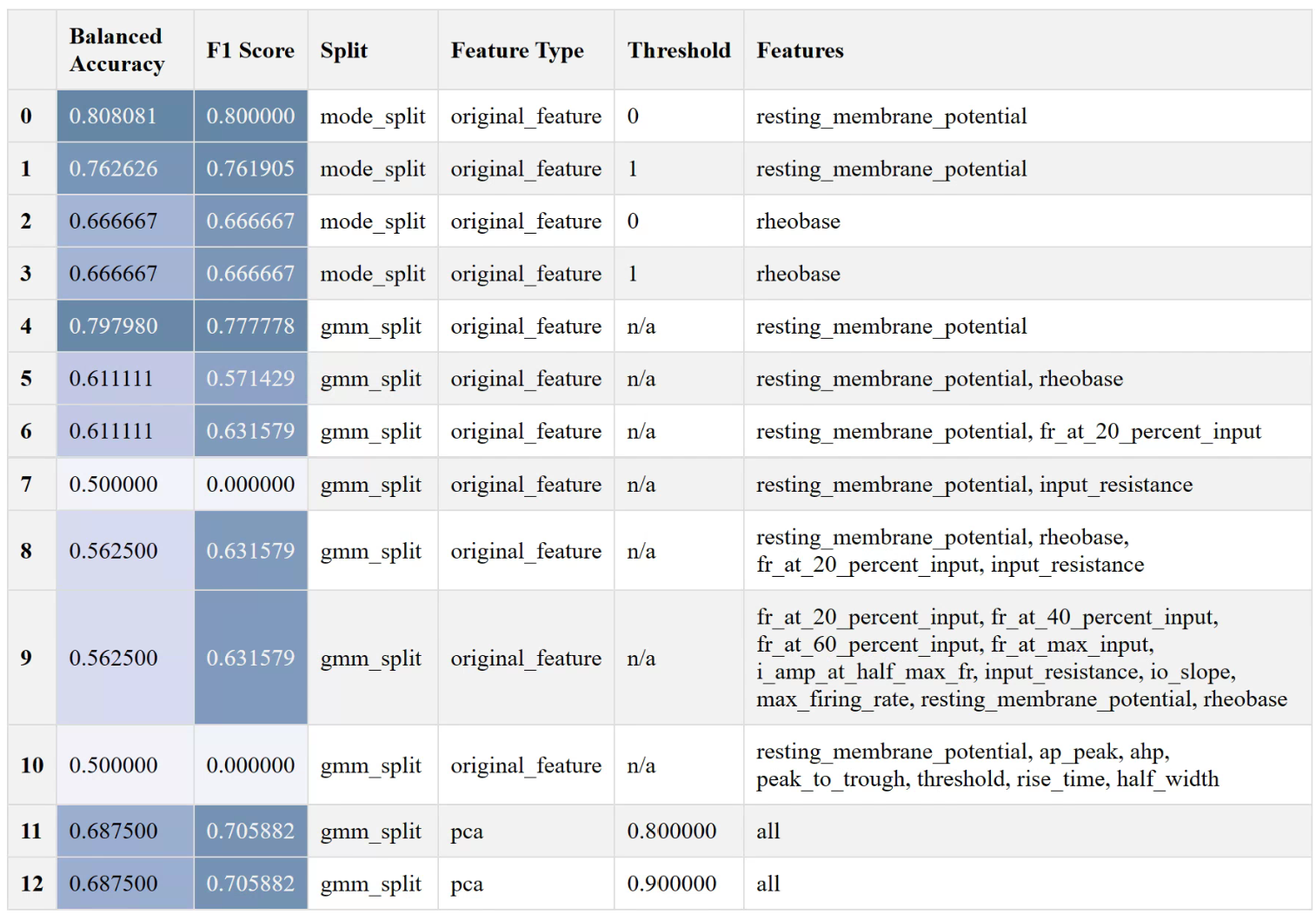

**Supplemental Table 1:** **Performance of Gaussian naïve Bayes model on predicting BnST vs NAcc BLA projecting cells in CUS mice.** Mode split indicates separation of clusters by mode of control cells, GMM split indicates separation of clusters using a GMM. Threshold in mode split represents the mode plus the standard deviation multiplier. Threshold in PCA represents the cumulative explained variance cutoff for the selection of principal components to be included for clustering.

| **Description** | **Figure** | **Stat test** | **Stats report** |
| --- | --- | --- | --- |
| Fr. 20% input | Figure 2C, Supp. Figure 3A | Mann–Whitney U & BH correction | U=36.0, p=0.190, Nbnst=9, Nnacc=6, |
| Fr. 40% input | Figure 2C, Supp. Figure 3A | Mann–Whitney U & BH correction | U=42.0, p=0.170, Nbnst=9, Nnacc=6, |
| Fr. 60% input | Figure 2C, Supp. Figure 3A | Mann–Whitney U & BH correction | U=40.0, p=0.190, Nbnst=9, Nnacc=6, |
| Fr. Max input | Figure 2C, Supp. Figure 3A | Mann–Whitney U & BH correction | U=22.5, p=0.636, Nbnst=9, Nnacc=6, |
| Current Inj. at Half Max Fr | Supp. Figure 3A | Mann–Whitney U & BH correction | U=6.5, p=0.059, Nbnst=9, Nnacc=6, |
| Input Resistance | Supp. Figure 3A | Mann–Whitney U & BH correction | U=68.0, p=0.165, Nbnst=10, Nnacc=9, |
| IO slope | Supp. Figure 3A | Mann–Whitney U & BH correction | U=47.0, p=0.059, Nbnst=9, Nnacc=6, |
| Max Fr. Rate | Supp. Figure 3A | Mann–Whitney U & BH correction | U=38.5, p=0.214, Nbnst=9, Nnacc=6, |
| Resting Membrane Potential | Figure 2D, Supp. Figure 3A | Mann–Whitney U & BH correction | U=91.5, p=0.016, Nbnst=11, Nnacc=9, |
| Rheobase | Supp. Figure 3A | Mann–Whitney U & BH correction | U=14.5, p=0.190, Nbnst=9, Nnacc=6, |
| AP amplitude | Supp. Figure 3A | Mann–Whitney U & BH correction | U=21.0, p=0.093, Nbnst=11, Nnacc=8, |
| AHP amplitude | Supp. Figure 3A | Mann–Whitney U & BH correction | U=33.0, p=0.395, Nbnst=11, Nnacc=8, |
| AP peak to trough | Supp. Figure 3A | Mann–Whitney U & BH correction | U=70.0, p=0.070, Nbnst=11, Nnacc=8, |
| AP threshold | Supp. Figure 3A | Mann–Whitney U & BH correction | U=64.0, p=0.130, Nbnst=11, Nnacc=8, |
| AP rise time | Figure 2B, Supp. Figure 3A | Mann–Whitney U & BH correction | U=77.0, p=0.039, Nbnst=11, Nnacc=8, |
| AP half width | Figure 2B, Supp. Figure 3A | Mann–Whitney U & BH correction | U=74.5, p=0.039, Nbnst=11, Nnacc=8, |
| Resting Membrane Potential 4 groups | Figure 2E | Kruskal-Wallis test | H(4, n=48)=33.84, p<0.0001 |
| cus- vs. nacc | Figure 2E | Dunn's multiple comparisons test | z=0.320, p>0.9999, Ncus-=11, Nnacc=9 |
| cus+ vs. bnst | Figure 2E | Dunn's multiple comparisons test | z=1.695, p=0.36, Ncus+=17, Nbnst=11 |
| nacc vs. cus+ | Figure 2E | Dunn's multiple comparisons test | z=4.379, p<0.0001, Nnacc=9, Ncus+=17 |
| cus- vs. bnst | Figure 2E | Dunn's multiple comparisons test | z=3.032, p=0.0097, Ncus-=11, Nbnst=11 |

**Stats Table 2.** Highlighted rows: p < 0.05

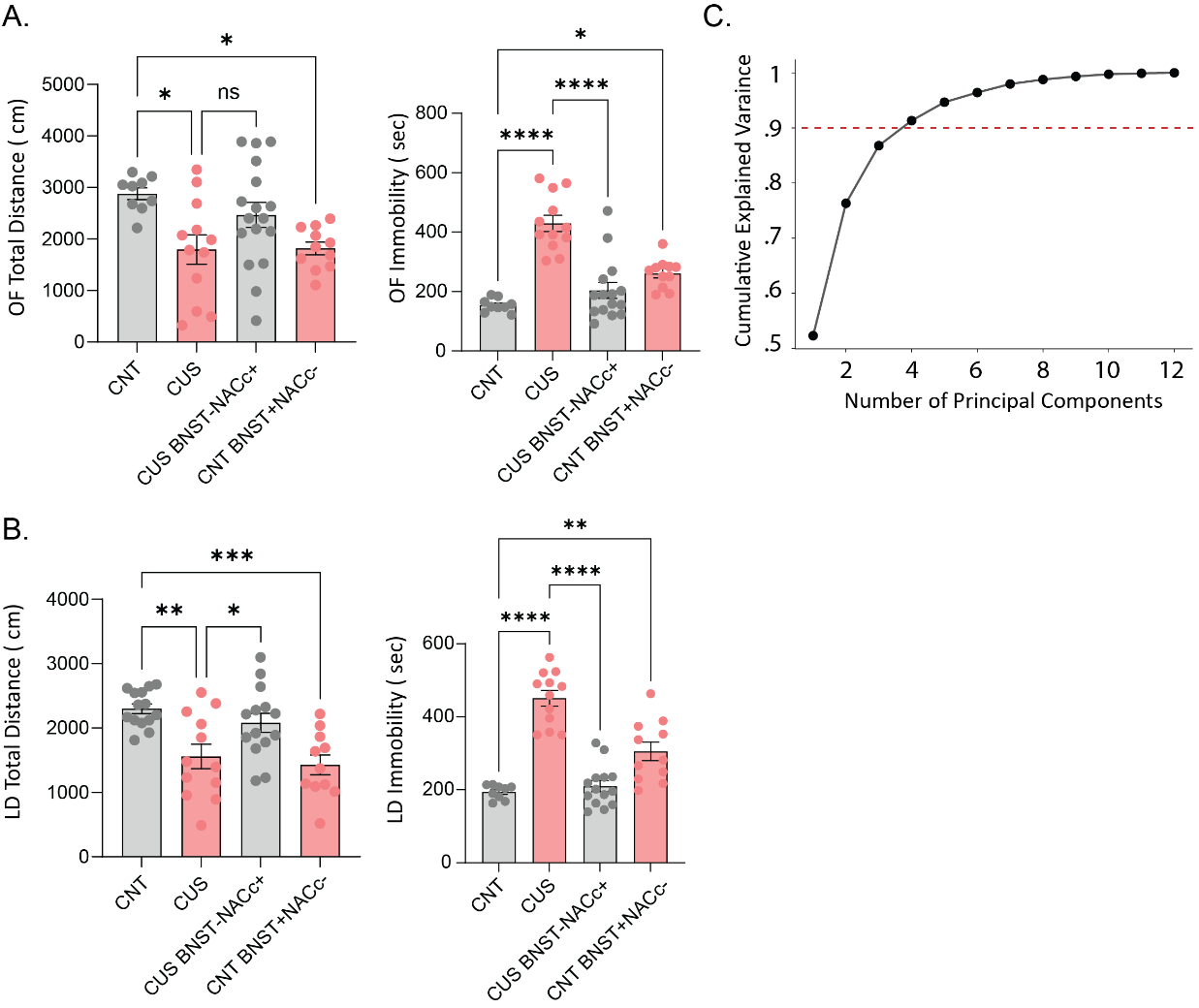

**Supplemental Figure 4: Activation of BLA-BnST pathway and inactivation of BLA-NAcc pathway mimics the behavioral effects of CUS on activity levels and freezing behavior.** CUS and CNT BnST+NAcc- exhibit a decrease in the total distance traveled and an increase in immobility in **A**) open field and **B**) light dark box. **C)** Cumulative explained variance ratio plotted against number of principal components with the threshold (red dotted line) for selection of principal components for clustering in figure 3F (n = 3 principal components selected).

| **Description** | **Figure** | **Stat test** | **Stats report** |
| --- | --- | --- | --- |
| OF center distance | Figure 3C | one-way ANOVA | F(3, 45)=1.587, p=0.2058 |
| CNT vs. CUS | Figure 3C | Sidak test | t=1.907, p=0.177, N1= 9, N2=12 |
| CUS vs. CUS BNST-NACc+ | Figure 3C | Sidak test | t=1.699, p=0.2618, N1= 12, N2=17 |
| CNT vs. CNT BNST+NACc- | Figure 3C | Sidak test | t=1.263, p=0.5128, N1= 9, N2=11 |
| OF center time | Figure 3C | one-way ANOVA | F(3, 45)=0.9375, p=0.4305 |
| CNT vs. CUS | Figure 3C | Sidak test | t=1.066, p=0.6456, N1= 9, N2=12 |
| CUS vs. CUS BNST-NACc+ | Figure 3C | Sidak test | t=1.543, p=0.3413, N1= 12, N2=17 |
| CNT vs. CNT BNST+NACc- | Figure 3C | Sidak test | t=0.2112, p=0.9954, N1= 9, N2=11 |
| OF center entries | Figure 3C | one-way ANOVA | F(3, 45)=1.787, p=0.1632 |
| CNT vs. CUS | Figure 3C | Sidak test | t=1.926, p=0.1706, N1= 9, N2=12 |
| CUS vs. CUS BNST-NACc+ | Figure 3C | Sidak test | t=1.646, p=0.2871, N1= 12, N2=17 |
| CNT vs. CNT BNST+NACc- | Figure 3C | Sidak test | t=1.598, p=0.3114, N1= 9, N2=11 |
| OF total distance | Supp. Figure 4A | one-way ANOVA | F(3, 45)=4.486, p=0.0077 |
| CNT vs. CUS | Supp. Figure 4A | Sidak test | t=3.026, p=0.0122, N1= 9, N2=12 |
| CUS vs. CUS BNST-NACc+ | Supp. Figure 4A | Sidak test | t=2.189, p=0.0982, N1= 12, N2=17 |
| CNT vs. CNT BNST+NACc- | Supp. Figure 4A | Sidak test | t=2.909, p=0.0167, N1= 9, N2=11 |
| OF total immobility | Supp. Figure 4A | one-way ANOVA | F(3, 43)=25.60, p<0.0001 |
| CNT vs. CUS | Supp. Figure 4A | Sidak test | t=7.797, p=<0.0001, N1= 9, N2=12 |
| CUS vs. CUS BNST-NACc+ | Supp. Figure 4A | Sidak test | t=7.291, p=<0.0001, N1= 12, N2=15 |
| CNT vs. CNT BNST+NACc- | Supp. Figure 4A | Sidak test | t=2.957, p=0.015, N1= 9, N2=11 |
| LD light distance | Figure 3D | one-way ANOVA | F(3, 47)=8.758, p=0.0001 |
| CNT vs. CUS | Figure 3D | Sidak test | t=3.49, p=0.0032, N1= 14, N2=12 |
| CUS vs. CUS BNST-NACc+ | Figure 3D | Sidak test | t=1.803, p=0.2157, N1= 12, N2=14 |
| CNT vs. CNT BNST+NACc- | Figure 3D | Sidak test | t=4.77, p=<0.0001, N1= 14, N2=11 |
| LD light time | Figure 3D | one-way ANOVA | F(3, 47)=8.453, p=0.0001 |
| CNT vs. CUS | Figure 3D | Sidak test | t=3.061, p=0.0109, N1= 14, N2=12 |
| CUS vs. CUS BNST-NACc+ | Figure 3D | Sidak test | t=0.8733, p=0.7696, N1= 12, N2=14 |
| CNT vs. CNT BNST+NACc- | Figure 3D | Sidak test | t=4.932, p=<0.0001, N1= 14, N2=11 |
| LD light entries | Figure 3D | one-way ANOVA | F(3, 47)=6.319, p=0.0011 |
| CNT vs. CUS | Figure 3D | Sidak test | t=3.399, p=0.0042, N1= 14, N2=12 |
| CUS vs. CUS BNST-NACc+ | Figure 3D | Sidak test | t=2.567, p=0.0399, N1= 12, N2=14 |
| CNT vs. CNT BNST+NACc- | Figure 3D | Sidak test | t=3.457, p=0.0035, N1= 14, N2=11 |
| LD total distance | Supp. Figure 4B | one-way ANOVA | F(3, 47)=8.325, p=0.0002 |
| CNT vs. CUS | Supp. Figure 4B | Sidak test | t=3.689, p=0.0017, N1= 14, N2=12 |
| CUS vs. CUS BNST-NACc+ | Supp. Figure 4B | Sidak test | t=2.606, p=0.0363, N1= 12, N2=14 |
| CNT vs. CNT BNST+NACc- | Supp. Figure 4B | Sidak test | t=4.231, p=0.0003, N1= 14, N2=11 |
| LD total immobility | Supp. Figure 4B | one-way ANOVA | F(3, 42)=39.08, p<0.0001 |
| CNT vs. CUS | Supp. Figure 4B | Sidak test | t=9.038, p=<0.0001, N1= 9, N2=12 |
| CUS vs. CUS BNST-NACc+ | Supp. Figure 4B | Sidak test | t=9.53, p=<0.0001, N1= 12, N2=14 |
| CNT vs. CNT BNST+NACc- | Supp. Figure 4B | Sidak test | t=3.834, p=0.0012, N1= 9, N2=11 |
| TST latency to immobility | Figure 3E | one-way ANOVA | F(3, 39)=26.40, p<0.0001 |
| CNT vs. CUS | Figure 3E | Sidak test | t=7.529, p=<0.0001, N1= 9, N2=8 |
| CUS vs. CUS BNST-NACc+ | Figure 3E | Sidak test | t=5.906, p=<0.0001, N1= 8, N2=15 |
| CNT vs. CNT BNST+NACc- | Figure 3E | Sidak test | t=6.556, p=<0.0001, N1= 9, N2=11 |
| TST total immobility | Figure 3E | one-way ANOVA | F(3, 39)=39.68, p<0.0001 |
| CNT vs. CUS | Figure 3E | Sidak test | t=7.342, p=<0.0001, N1= 9, N2=8 |
| CUS vs. CUS BNST-NACc+ | Figure 3E | Sidak test | t=9.154, p=<0.0001, N1= 8, N2=15 |
| CNT vs. CNT BNST+NACc- | Figure 3E | Sidak test | t=5.738, p=<0.0001, N1= 9, N2=11 |

**Stats Table 3.** Highlighted rows: p < 0.05

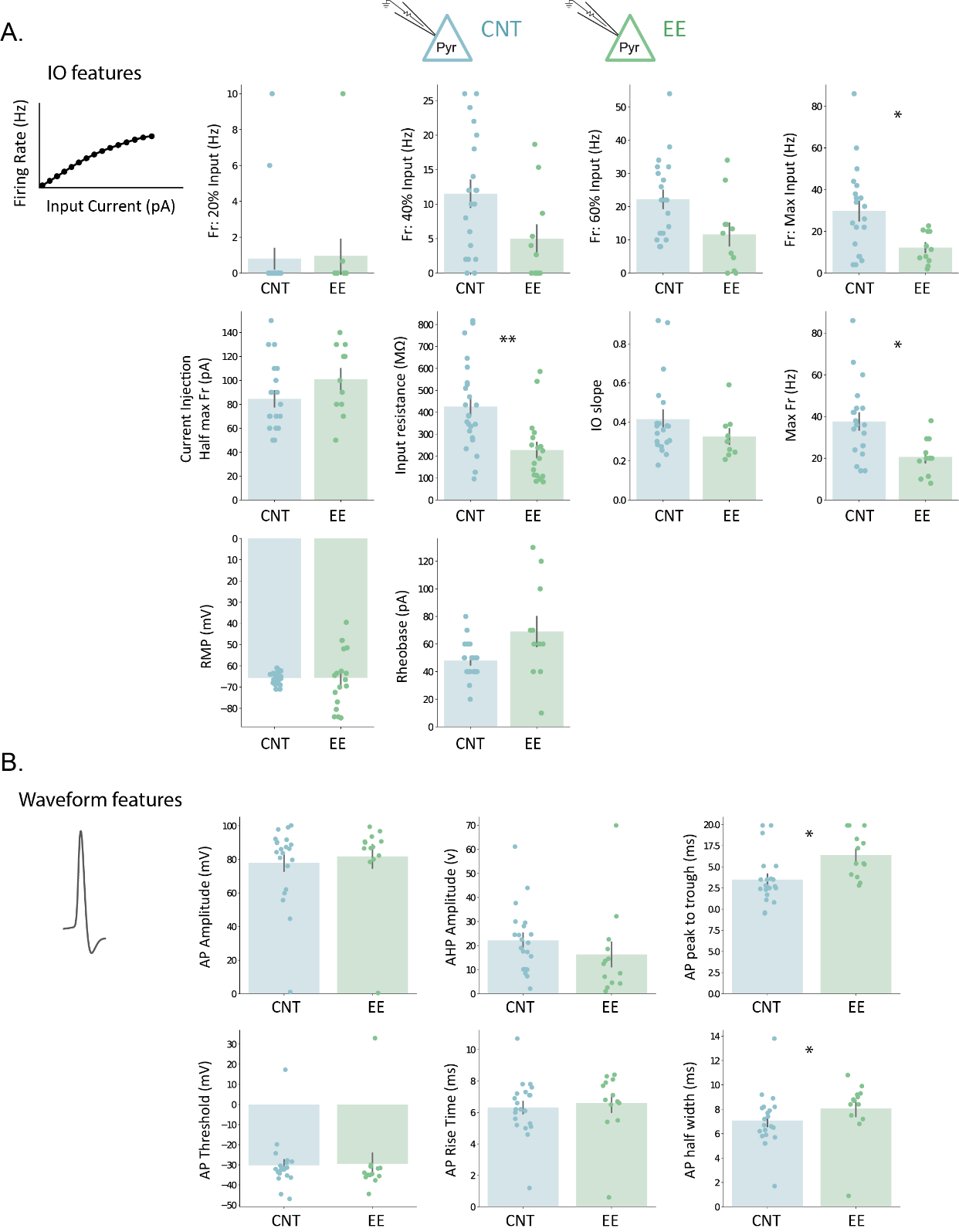

**Supplemental Figure 5:** **Effect of enriched environment on BLA principal cell properties.** **A)** Bar plots with individual points for input-output (IO) properties/features. **B)** Bar plots with individual points for action potential waveform properties/features.

| **Description** | **Figure** | **Stat test** | **Stats report** |
| --- | --- | --- | --- |
| Fr. 20% input | Figure 4C, Supp. Figure 5A | Mann–Whitney U & BH correction | U=101.5, p=0.634, Ncontrol=20, Nee=11, |
| Fr. 40% input | Figure 4C, Supp. Figure 5A | Mann–Whitney U & BH correction | U=161.5, p=0.077, Ncontrol=20, Nee=11, |
| Fr. 60% input | Figure 4C, Supp. Figure 5A | Mann–Whitney U & BH correction | U=160.5, p=0.077, Ncontrol=20, Nee=11, |
| Fr. Max input | Figure 4C, Supp. Figure 5A | Mann–Whitney U & BH correction | U=174.5, p=0.027, Ncontrol=20, Nee=11, |
| Current Inj. at Half Max Fr | Supp. Figure 5A | Mann–Whitney U & BH correction | U=73.5, p=0.193, Ncontrol=20, Nee=11, |
| Input Resistance | Figure 4D, Supp. Figure 5A | Mann–Whitney U & BH correction | U=351.0, p=0.006, Ncontrol=24, Nee=18, |
| IO slope | Supp. Figure 5A | Mann–Whitney U & BH correction | U=117.0, p=0.264, Ncontrol=20, Nee=9, |
| Max Fr. Rate | Supp. Figure 5A | Mann–Whitney U & BH correction | U=178.5, p=0.025, Ncontrol=20, Nee=11, |
| Resting Membrane Potential | Supp. Figure 5A | Mann–Whitney U & BH correction | U=226.0, p=0.809, Ncontrol=24, Nee=18, |
| Rheobase | Supp. Figure 5A | Mann–Whitney U & BH correction | U=62.5, p=0.080, Ncontrol=20, Nee=11, |
| AP amplitude | Supp. Figure 5B | Mann–Whitney U & BH correction | U=112.0, p=0.395, Ncontrol=21, Nee=13, |
| AHP amplitude | Supp. Figure 5B | Mann–Whitney U & BH correction | U=188.0, p=0.141, Ncontrol=21, Nee=13, |
| AP peak to trough | Figure 4B, Supp. Figure 5B | Mann–Whitney U & BH correction | U=49.5, p=0.013, Ncontrol=21, Nee=13, |
| AP threshold | Supp. Figure 5B | Mann–Whitney U & BH correction | U=168.0, p=0.326, Ncontrol=21, Nee=13, |
| AP rise time | Supp. Figure 5B | Mann–Whitney U & BH correction | U=102.5, p=0.326, Ncontrol=21, Nee=13, |
| AP half width | Figure 4B, Supp. Figure 5B | Mann–Whitney U & BH correction | U=67.0, p=0.043, Ncontrol=21, Nee=13, |
| input-output properties, Control-clusters | Figure 4E | Permutation-based MANOVA | pseudo-F(1, 18) = 0.345, p < 0.886 |
| wave properties, Control-clusters | Figure 4E | Permutation-based MANOVA | pseudo-F(1, 19) = 1.760, p < 0.166 |
| Fr. 20% input, Control-clusters | Figure 4E | Mann–Whitney U & BH correction | U=51.0, p=0.969, Ncontrol+=9, Ncontrol-=11, |
| Fr. 40% input, Control-clusters | Figure 4E | Mann–Whitney U & BH correction | U=41.5, p=0.969, Ncontrol+=9, Ncontrol-=11, |
| Fr. 60% input, Control-clusters | Figure 4E | Mann–Whitney U & BH correction | U=43.0, p=0.969, Ncontrol+=9, Ncontrol-=11, |
| Fr. Max input, Control-clusters | Figure 4E | Mann–Whitney U & BH correction | U=55.5, p=0.969, Ncontrol+=9, Ncontrol-=11, |
| Current Inj. at Half Max Fr, Control-clusters | Figure 4E | Mann–Whitney U & BH correction | U=51.5, p=0.969, Ncontrol+=9, Ncontrol-=11, |
| Input Resistance, Control-clusters | Figure 4E | Mann–Whitney U & BH correction | U=51.0, p=0.969, Ncontrol+=9, Ncontrol-=15, |
| IO slope, Control-clusters | Figure 4E | Mann–Whitney U & BH correction | U=45.0, p=0.969, Ncontrol+=9, Ncontrol-=11, |
| Max Fr. Rate, Control-clusters | Figure 4E | Mann–Whitney U & BH correction | U=44.5, p=0.969, Ncontrol+=9, Ncontrol-=11, |
| Rheobase, Control-clusters | Figure 4E | Mann–Whitney U & BH correction | U=48.5, p=0.969, Ncontrol+=9, Ncontrol-=11, |
| input-output properties, EE-clusters | Figure 4E | Permutation-based MANOVA | pseudo-F(1, 7) = 1.767, p < 0.110 |
| wave properties, EE-clusters | Figure 4E | Permutation-based MANOVA | pseudo-F(1, 11) = 0.668, p < 0.856 |
| Fr. 20% input, EE-clusters | Figure 4E | Mann–Whitney U & BH correction | U=20.0, p=0.279, Nee+=6, Nee-=5, |
| Fr. 40% input, EE-clusters | Figure 4E | Mann–Whitney U & BH correction | U=30.0, p=0.018, Nee+=6, Nee-=5, |
| Fr. 60% input, EE-clusters | Figure 4E | Mann–Whitney U & BH correction | U=27.0, p=0.058, Nee+=6, Nee-=5, |
| Fr. Max input, EE-clusters | Figure 4E | Mann–Whitney U & BH correction | U=6.0, p=0.171, Nee+=6, Nee-=5, |
| Current Inj. at Half Max Fr, EE-clusters | Figure 4E | Mann–Whitney U & BH correction | U=9.5, p=0.358, Nee+=6, Nee-=5, |
| Input Resistance, EE-clusters | Figure 4E | Mann–Whitney U & BH correction | U=73.0, p=0.010, Nee+=8, Nee-=10, |
| IO slope, EE-clusters | Figure 4E | Mann–Whitney U & BH correction | U=18.0, p=0.048, Nee+=6, Nee-=3, |
| Max Fr. Rate, EE-clusters | Figure 4E | Mann–Whitney U & BH correction | U=21.0, p=0.344, Nee+=6, Nee-=5, |
| Rheobase, EE-clusters | Figure 4E | Mann–Whitney U & BH correction | U=0.0, p=0.018, Nee+=6, Nee-=5, |

**Stats Table 4.** Highlighted rows: p < 0.05

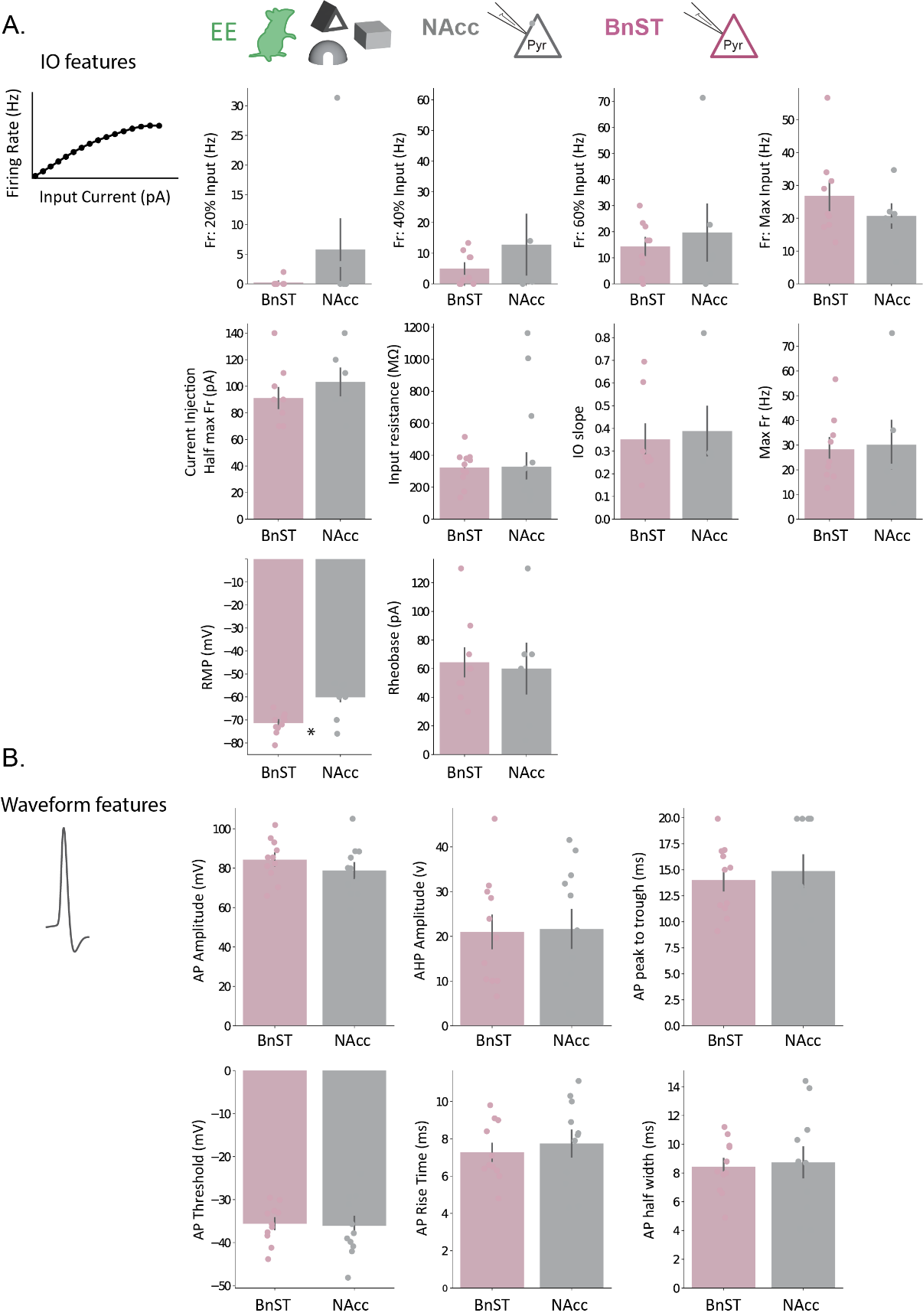

**Supplemental Figure 6:** **Projection-specific effect of enriched environment on BLA principal cell properties. A)** Bar plots with individual points for input-output (IO) properties/features of BLA-BnST and BLA-NAcc populations. **B)** Bar plots with individual points for action potential waveform properties/features from BLA-BnST and BLA-NAcc neurons.

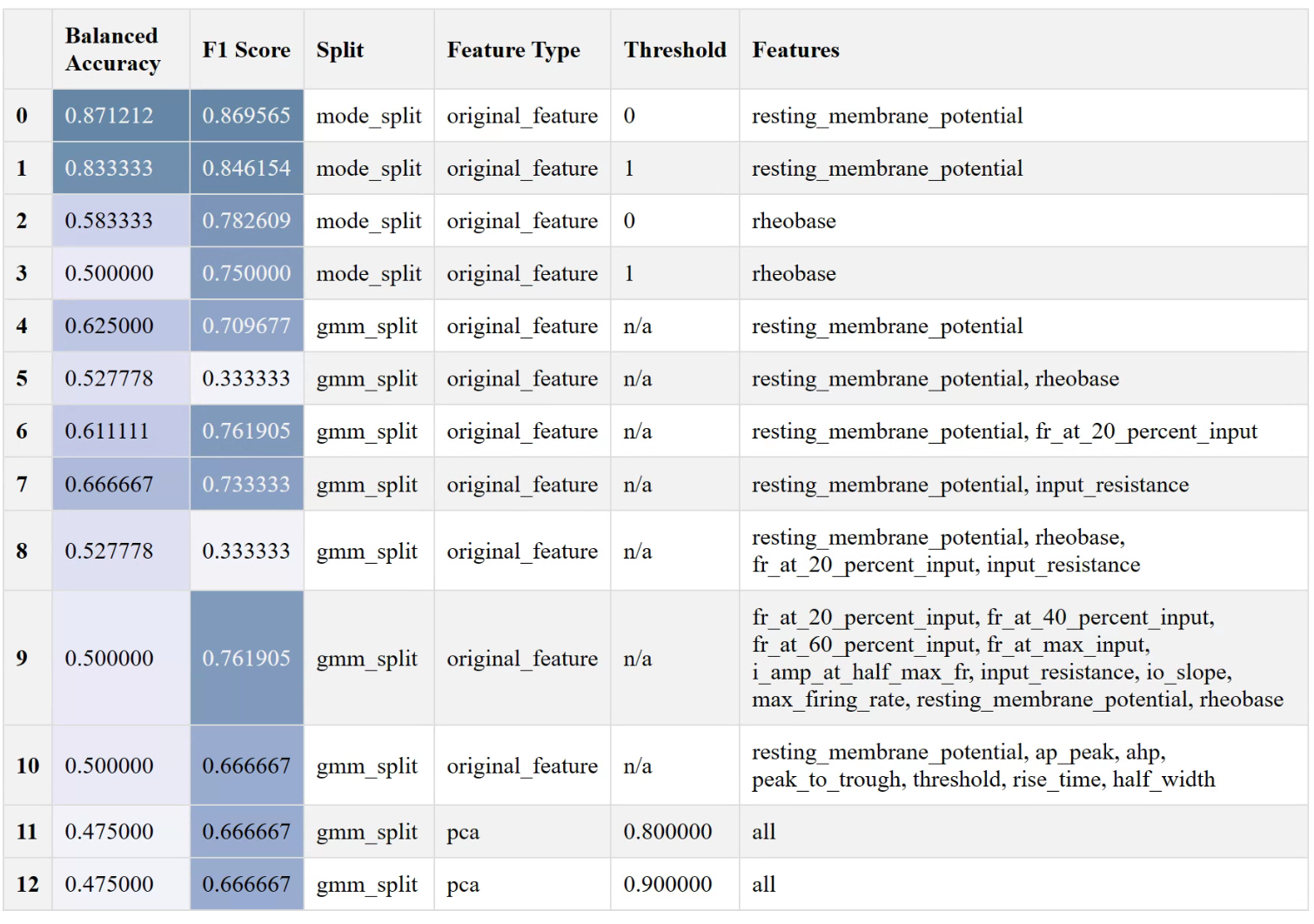

**Supplemental Table 2:** **Performance of Gaussian naïve Bayes model on predicting BnST vs NAcc BLA projecting cells in EE mice.** Mode split indicates separation of clusters by mode of control cells, GMM split indicates separation of clusters using a GMM. Threshold in mode split represents the mode + the standard deviation multiplier. Threshold in PCA represents the cumulative explained variance cutoff for the selection of principal components to be included for clustering.

| **Description** | **Figure** | **Stat test** | **Stats report** |
| --- | --- | --- | --- |
| Fr. 20% input | Figure 5A, Supp. Figure 6A | Mann–Whitney U & BH correction | U=20.0, p=0.840, Nbnst=9, Nnacc=6, |
| Fr. 40% input | Figure 5A, Supp. Figure 6A | Mann–Whitney U & BH correction | U=26.5, p=1.000, Nbnst=9, Nnacc=6, |
| Fr. 60% input | Figure 5A, Supp. Figure 6A | Mann–Whitney U & BH correction | U=29.0, p=1.000, Nbnst=9, Nnacc=6, |
| Fr. Max input | Figure 5A, Supp. Figure 6A | Mann–Whitney U & BH correction | U=30.5, p=1.000, Nbnst=9, Nnacc=6, |
| Current Inj. at Half Max Fr | Supp. Figure 6A | Mann–Whitney U & BH correction | U=18.5, p=0.840, Nbnst=9, Nnacc=6, |
| Input Resistance | Supp. Figure 6A | Mann–Whitney U & BH correction | U=87.0, p=0.840, Nbnst=11, Nnacc=12, |
| IO slope | Supp. Figure 6A | Mann–Whitney U & BH correction | U=15.0, p=1.000, Nbnst=8, Nnacc=5, |
| Max Fr. Rate | Supp. Figure 6A | Mann–Whitney U & BH correction | U=27.5, p=1.000, Nbnst=9, Nnacc=6, |
| Resting Membrane Potential | Supp. Figure 6A | Mann–Whitney U & BH correction | U=15.0, p=0.018, Nbnst=11, Nnacc=12, |
| Rheobase | Supp. Figure 6A | Mann–Whitney U & BH correction | U=27.5, p=1.000, Nbnst=9, Nnacc=6, |
| AP amplitude | Supp. Figure 6A | Mann–Whitney U & BH correction | U=79.0, p=0.922, Nbnst=11, Nnacc=11, |
| AHP amplitude | Supp. Figure 6A | Mann–Whitney U & BH correction | U=57.0, p=0.922, Nbnst=11, Nnacc=11, |
| AP peak to trough | Supp. Figure 6A | Mann–Whitney U & BH correction | U=54.5, p=0.922, Nbnst=11, Nnacc=11, |
| AP threshold | Supp. Figure 6A | Mann–Whitney U & BH correction | U=63.0, p=0.922, Nbnst=11, Nnacc=11, |
| AP rise time | Supp. Figure 6A | Mann–Whitney U & BH correction | U=56.0, p=0.922, Nbnst=11, Nnacc=11, |
| AP half width | Supp. Figure 6A | Mann–Whitney U & BH correction | U=58.5, p=0.922, Nbnst=11, Nnacc=11, |
| Resting Membrane Potential 4 groups | Figure 5D | Kruskal-Wallis test | H(4, n=41)=25.23, p<0.0001 |
| ee- vs. bnst | Figure 5D | Dunn's multiple comparisons test | z=0.664, p=>0.9999, N1= 10, N2=11 |
| ee+ vs. nacc | Figure 5D | Dunn's multiple comparisons test | z=1.045, p=>0.9999, N1= 8, N2=12 |
| bnst vs. ee+ | Figure 5D | Dunn's multiple comparisons test | z=3.592, p=0.0013, N1= 11, N2=8 |
| ee- vs. nacc | Figure 5D | Dunn's multiple comparisons test | z=3.462, p=0.0021, N1= 10, N2=12 |

**Stats Table 5.** Highlighted rows: p < 0.05

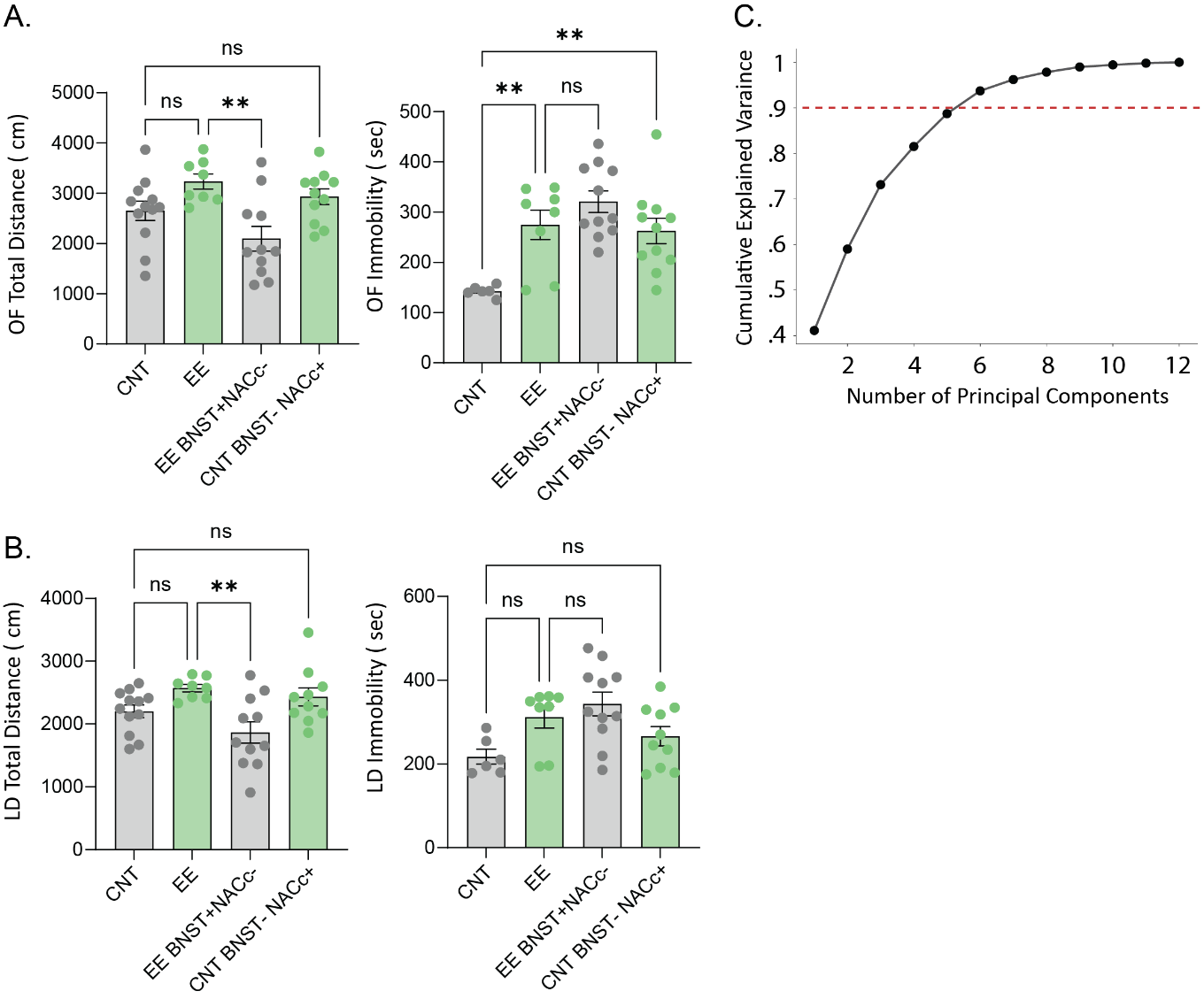

**Supplemental Figure 7**: **Manipulation of BLA-NAcc and BLA-BnST circuits alters activity levels and freezing behavior following enriched environment exposure**. The total distance traveled and immobility in **A**) open field and **B**) light dark box. **C)** Cumulative explained variance ratio plotted against number of principal components with the threshold (red dotted line) for selection of principal components for clustering in figure 6F (n = 5 principal components selected).

| **Description** | **Figure** | **Stat test** | **Stats report** |
| --- | --- | --- | --- |
| OF center distance | Figure 6C | one-way ANOVA | F(3, 38)=6.851, p=0.0008 |
| CNT vs. EE | Figure 6C | Sidak test | t=2.171, p=0.1048, N1= 12, N2=8 |
| EE vs. EE BNST+NACc- | Figure 6C | Sidak test | t=3.217, p=0.0079, N1= 8, N2=11 |
| CNT vs. CNT BNST- NACc+ | Figure 6C | Sidak test | t=2.855, p=0.0207, N1= 12, N2=11 |
| OF center time | Figure 6C | one-way ANOVA | F(3, 38)=4.746, p=0.0066 |
| CNT vs. EE | Figure 6C | Sidak test | t=0.8406, p=0.7902, N1= 12, N2=8 |
| EE vs. EE BNST+NACc- | Figure 6C | Sidak test | t=1.461, p=0.391, N1= 8, N2=11 |
| CNT vs. CNT BNST- NACc+ | Figure 6C | Sidak test | t=2.912, p=0.0178, N1= 12, N2=11 |
| OF center entries | Figure 6C | one-way ANOVA | F(3, 38)=7.203, p=0.0006 |
| CNT vs. EE | Figure 6C | Sidak test | t=2.721, p=0.029, N1= 12, N2=8 |
| EE vs. EE BNST+NACc- | Figure 6C | Sidak test | t=3.877, p=0.0012, N1= 8, N2=11 |
| CNT vs. CNT BNST- NACc+ | Figure 6C | Sidak test | t=2.4, p=0.0628, N1= 12, N2=11 |
| OF total distance | Supp. Figure 7A | one-way ANOVA | F(3, 38)=5.737, p=0.0024 |
| CNT vs. EE | Supp. Figure 7A | Sidak test | t=2.017, p=0.1448, N1= 12, N2=8 |
| EE vs. EE BNST+NACc- | Supp. Figure 7A | Sidak test | t=3.864, p=0.0013, N1= 8, N2=11 |
| CNT vs. CNT BNST- NACc+ | Supp. Figure 7A | Sidak test | t=1.067, p=0.6464, N1= 12, N2=11 |
| OF total immobility | Supp. Figure 7A | one-way ANOVA | F(3, 32)=7.808, p=0.0005 |
| CNT vs. EE | Supp. Figure 7A | Sidak test | t=3.346, p=0.0063, N1= 6, N2=8 |
| EE vs. EE BNST+NACc- | Supp. Figure 7A | Sidak test | t=1.365, p=0.452, N1= 8, N2=11 |
| CNT vs. CNT BNST- NACc+ | Supp. Figure 7A | Sidak test | t=3.234, p=0.0085, N1= 6, N2=11 |
| LD light distance | Figure 6D | one-way ANOVA | F(3, 37)=3.782, p=0.0184 |
| CNT vs. EE | Figure 6D | Sidak test | t=2.317, p=0.0763, N1= 12, N2=8 |
| EE vs. EE BNST+NACc- | Figure 6D | Sidak test | t=3.217, p=0.008, N1= 8, N2=11 |
| CNT vs. CNT BNST- NACc+ | Figure 6D | Sidak test | t=0.9965, p=0.6931, N1= 12, N2=10 |
| LD light time | Figure 6D | one-way ANOVA | F(3, 37)=3.057, p=0.0402 |
| CNT vs. EE | Figure 6D | Sidak test | t=1.78, p=0.2295, N1= 12, N2=8 |
| EE vs. EE BNST+NACc- | Figure 6D | Sidak test | t=3.012, p=0.0139, N1= 8, N2=11 |
| CNT vs. CNT BNST- NACc+ | Figure 6D | Sidak test | t=0.269, p=0.9907, N1= 12, N2=10 |
| LD light entries | Figure 6D | one-way ANOVA | F(3, 37)=6.953, p=0.0008 |
| CNT vs. EE | Figure 6D | Sidak test | t=3.967, p=0.001, N1= 12, N2=8 |
| EE vs. EE BNST+NACc- | Figure 6D | Sidak test | t=4.099, p=0.0007, N1= 8, N2=11 |
| CNT vs. CNT BNST- NACc+ | Figure 6D | Sidak test | t=1.528, p=0.3529, N1= 12, N2=10 |
| LD total distance | Supp. Figure 7B | one-way ANOVA | F(3, 37)=5.232, p=0.0041 |
| CNT vs. EE | Supp. Figure 7B | Sidak test | t=1.908, p=0.1804, N1= 12, N2=8 |
| EE vs. EE BNST+NACc- | Supp. Figure 7B | Sidak test | t=3.596, p=0.0028, N1= 8, N2=11 |
| CNT vs. CNT BNST- NACc+ | Supp. Figure 7B | Sidak test | t=1.257, p=0.5191, N1= 12, N2=10 |
| LD total immobility | Supp. Figure 7B | one-way ANOVA | F(3, 31)=4.098, p=0.0147 |
| CNT vs. EE | Supp. Figure 7B | Sidak test | t=2.271, p=0.088, N1= 6, N2=8 |
| EE vs. EE BNST+NACc- | Supp. Figure 7B | Sidak test | t=0.9001, p=0.7559, N1= 8, N2=11 |
| CNT vs. CNT BNST- NACc+ | Supp. Figure 7B | Sidak test | t=1.231, p=0.5392, N1= 6, N2=10 |
| TST latency to immobility | Figure 6E | one-way ANOVA | F(3, 22)=3.083, p=0.0484 |
| CNT vs. EE | Figure 6E | Sidak test | t=0.05912, p=0.9999, N1= 9, N2=6 |
| EE vs. EE BNST+NACc- | Figure 6E | Sidak test | t=2.192, p=0.1132, N1= 6, N2=5 |
| CNT vs. CNT BNST- NACc+ | Figure 6E | Sidak test | t=0.8867, p=0.7672, N1= 9, N2=6 |
| TST total immobility | Figure 6E | one-way ANOVA | F(3, 22)=20.78, P<0.0001 |
| CNT vs. EE | Figure 6E | Sidak test | t=1.307, p=0.4967, N1= 9, N2=6 |
| EE vs. EE BNST+NACc- | Figure 6E | Sidak test | t=6.248, p=<0.0001, N1= 6, N2=5 |
| CNT vs. CNT BNST- NACc+ | Figure 6E | Sidak test | t=2.674, p=0.041, N1= 9, N2=6 |

**Stats Table 6.** Highlighted rows: p < 0.05

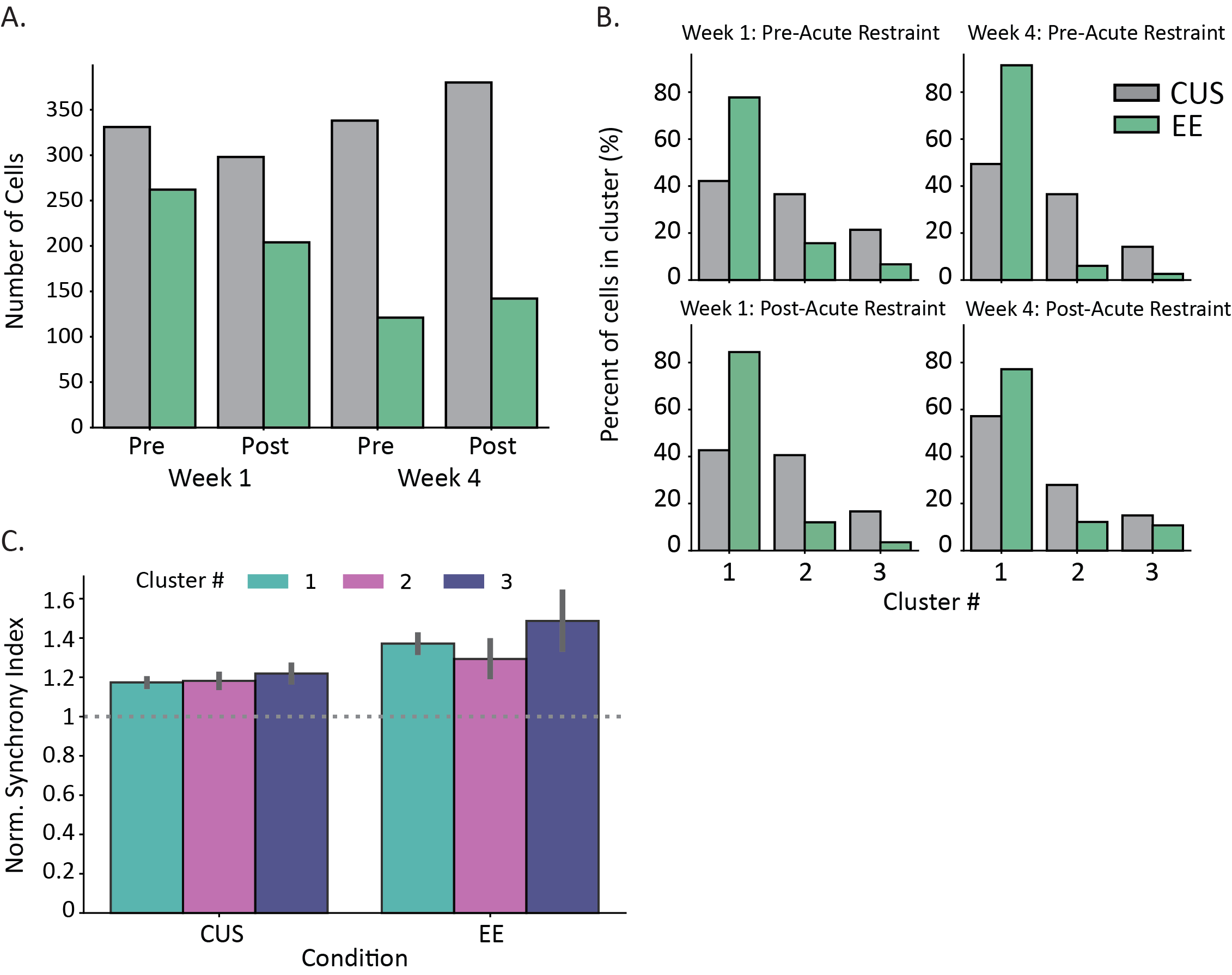

**Supplemental Figure 8:** **Characterization of imaged cells and calcium synchrony.** **A)** Grouped bar plot shows the impact of acute restraint on cell count from baseline (week 1) and condition (week 4) following CUS or EE. Chi-square tests response*week revealed no significant difference across conditions. **B)** Grouped bar plots show the impact of acute restraint on proportion of cells (across clusters) from baseline (week 1) and experience (week 4) following CUS or EE. Chi-square tests for each cluster (response*week) revealed no significant difference across conditions. **C)** Grouped bar plot by condition shows, across recordings, intra-cluster synchrony to be significantly higher than inter-cluster synchrony (Norm. Synch. Index > 1) for each cluster.

| **Description** | **Figure** | **Stat test** | **Stats report** |
| --- | --- | --- | --- |
| CUS: Pre/Post response; Height | Figure 6C/D | independent t-test | t=-5.814, p=<0.0001, N1=298, N2=380 |
| CUS: Pre/Post response; AUC | Figure 6C/D | independent t-test | t=-4.729, p=<0.0001, N1=298, N2=380 |
| CUS: Pre/Post response; Width | Figure 6C/D | independent t-test | t=0.411, p=0.6812, N1=298, N2=380 |
| CUS: Pre/Post response; Prominence | Figure 6C/D | independent t-test | t=-4.65, p=<0.0001, N1=298, N2=380 |
| CUS: Pre/Post response; Rise Time | Figure 6C/D | independent t-test | t=-1.918, p=0.0554, N1=298, N2=380 |
| CUS: Pre/Post response; Decay Time | Figure 6C/D | independent t-test | t=2.753, p=0.006, N1=298, N2=380 |
| CUS: Pre/Post response; Ca Events | Figure 6C/D | independent t-test | t=-7.142, p=<0.0001, N1=298, N2=380 |
| EE: Pre/Post response; Height | Figure 6C/D | independent t-test | t=1.854, p=0.0642, N1=204, N2=142 |
| EE: Pre/Post response; AUC | Figure 6C/D | independent t-test | t=1.762, p=0.0786, N1=204, N2=142 |
| EE: Pre/Post response; Width | Figure 6C/D | independent t-test | t=0.031, p=0.975, N1=204, N2=142 |
| EE: Pre/Post response; Prominence | Figure 6C/D | independent t-test | t=1.382, p=0.1675, N1=204, N2=142 |
| EE: Pre/Post response; Rise Time | Figure 6C/D | independent t-test | t=1.719, p=0.0861, N1=204, N2=142 |
| EE: Pre/Post response; Decay Time | Figure 6C/D | independent t-test | t=-3.347, p=0.0009, N1=204, N2=142 |
| EE: Pre/Post response; Ca Events | Figure 6C/D | independent t-test | t=-4.418, p=<0.0001, N1=204, N2=142 |
| CUS, Cluster 1: Pre/Post response; Height | Figure 6G/H | independent t-test | t=-5.255, p=<0.0001, N1=123, N2=211 |
| CUS, Cluster 1: Pre/Post response; AUC | Figure 6G/H | independent t-test | t=-6.363, p=<0.0001, N1=123, N2=211 |
| CUS, Cluster 1: Pre/Post response; Width | Figure 6G/H | independent t-test | t=-2.868, p=0.0043, N1=123, N2=211 |
| CUS, Cluster 1: Pre/Post response; Prominence | Figure 6G/H | independent t-test | t=-3.557, p=0.0004, N1=123, N2=211 |
| CUS, Cluster 1: Pre/Post response; Rise Time | Figure 6G/H | independent t-test | t=-4.056, p=<0.0001, N1=123, N2=211 |
| CUS, Cluster 1: Pre/Post response; Decay Time | Figure 6G/H | independent t-test | t=2.209, p=0.0276, N1=123, N2=211 |
| CUS, Cluster 1: Pre/Post response; Ca Events | Figure 6G/H | independent t-test | t=-4.12, p=<0.0001, N1=123, N2=211 |
| CUS, Cluster 2: Pre/Post response; Height | Figure 6G/H | independent t-test | t=-1.864, p=0.0629, N1=117, N2=103 |
| CUS, Cluster 2: Pre/Post response; AUC | Figure 6G/H | independent t-test | t=1.772, p=0.077, N1=117, N2=103 |
| CUS, Cluster 2: Pre/Post response; Width | Figure 6G/H | independent t-test | t=0.89, p=0.3739, N1=117, N2=103 |
| CUS, Cluster 2: Pre/Post response; Prominence | Figure 6G/H | independent t-test | t=-3.914, p=0.0001, N1=117, N2=103 |
| CUS, Cluster 2: Pre/Post response; Rise Time | Figure 6G/H | independent t-test | t=1.995, p=0.0466, N1=117, N2=103 |
| CUS, Cluster 2: Pre/Post response; Decay Time | Figure 6G/H | independent t-test | t=-1.654, p=0.0988, N1=117, N2=103 |
| CUS, Cluster 2: Pre/Post response; Ca Events | Figure 6G/H | independent t-test | t=-6.702, p=<0.0001, N1=117, N2=103 |
| CUS, Cluster 3: Pre/Post response; Height | Figure 6G/H | independent t-test | t=-2.63, p=0.0091, N1=48, N2=55 |
| CUS, Cluster 3: Pre/Post response; AUC | Figure 6G/H | independent t-test | t=-1.971, p=0.05, N1=48, N2=55 |
| CUS, Cluster 3: Pre/Post response; Width | Figure 6G/H | independent t-test | t=1.034, p=0.3025, N1=48, N2=55 |
| CUS, Cluster 3: Pre/Post response; Prominence | Figure 6G/H | independent t-test | t=-1.44, p=0.1514, N1=48, N2=55 |
| CUS, Cluster 3: Pre/Post response; Rise Time | Figure 6G/H | independent t-test | t=1.261, p=0.2088, N1=48, N2=55 |
| CUS, Cluster 3: Pre/Post response; Decay Time | Figure 6G/H | independent t-test | t=0.09, p=0.928, N1=48, N2=55 |
| CUS, Cluster 3: Pre/Post response; Ca Events | Figure 6G/H | independent t-test | t=-1.602, p=0.1107, N1=48, N2=55 |
| EE, Cluster 1: Pre/Post response; Height | Figure 6G/H | independent t-test | t=4.188, p=<0.0001, N1=169, N2=108 |
| EE, Cluster 1: Pre/Post response; AUC | Figure 6G/H | independent t-test | t=3.143, p=0.0018, N1=169, N2=108 |
| EE, Cluster 1: Pre/Post response; Width | Figure 6G/H | independent t-test | t=0.911, p=0.3625, N1=169, N2=108 |
| EE, Cluster 1: Pre/Post response; Prominence | Figure 6G/H | independent t-test | t=3.033, p=0.0025, N1=169, N2=108 |
| EE, Cluster 1: Pre/Post response; Rise Time | Figure 6G/H | independent t-test | t=2.703, p=0.0071, N1=169, N2=108 |
| EE, Cluster 1: Pre/Post response; Decay Time | Figure 6G/H | independent t-test | t=-3.492, p=0.0005, N1=169, N2=108 |
| EE, Cluster 1: Pre/Post response; Ca Events | Figure 6G/H | independent t-test | t=-2.155, p=0.0316, N1=169, N2=108 |
| EE, Cluster 2: Pre/Post response; Height | Figure 6G/H | independent t-test | t=-0.792, p=0.4303, N1=24, N2=17 |
| EE, Cluster 2: Pre/Post response; AUC | Figure 6G/H | independent t-test | t=2.774, p=0.0068, N1=24, N2=17 |
| EE, Cluster 2: Pre/Post response; Width | Figure 6G/H | independent t-test | t=4.375, p=<0.0001, N1=24, N2=17 |
| EE, Cluster 2: Pre/Post response; Prominence | Figure 6G/H | independent t-test | t=-0.774, p=0.4409, N1=24, N2=17 |
| EE, Cluster 2: Pre/Post response; Rise Time | Figure 6G/H | independent t-test | t=4.737, p=<0.0001, N1=24, N2=17 |
| EE, Cluster 2: Pre/Post response; Decay Time | Figure 6G/H | independent t-test | t=0.341, p=0.734, N1=24, N2=17 |
| EE, Cluster 2: Pre/Post response; Ca Events | Figure 6G/H | independent t-test | t=-1.695, p=0.0938, N1=24, N2=17 |
| EE, Cluster 3: Pre/Post response; Height | Figure 6G/H | independent t-test | t=3.72, p=0.0006, N1=7, N2=15 |
| EE, Cluster 3: Pre/Post response; AUC | Figure 6G/H | independent t-test | t=2.655, p=0.0113, N1=7, N2=15 |
| EE, Cluster 3: Pre/Post response; Width | Figure 6G/H | independent t-test | t=-1.294, p=0.203, N1=7, N2=15 |
| EE, Cluster 3: Pre/Post response; Prominence | Figure 6G/H | independent t-test | t=3.201, p=0.0027, N1=7, N2=15 |
| EE, Cluster 3: Pre/Post response; Rise Time | Figure 6G/H | independent t-test | t=-0.983, p=0.3316, N1=7, N2=15 |
| EE, Cluster 3: Pre/Post response; Decay Time | Figure 6G/H | independent t-test | t=-1.473, p=0.1486, N1=7, N2=15 |
| EE, Cluster 3: Pre/Post response; Ca Events | Figure 6G/H | independent t-test | t=-0.216, p=0.83, N1=7, N2=15 |
| CUS, Cluster 1: Pre/Post response; Synchrony | Figure 6K | independent t-test | t=3.26, p=0.0012, N1=117, N2=205 |
| CUS, Cluster 2: Pre/Post response; Synchrony | Figure 6K | independent t-test | t=9.115, p=<0.0001, N1=111, N2=97 |
| CUS, Cluster 3: Pre/Post response; Synchrony | Figure 6K | independent t-test | t=-5.02, p=<0.0001, N1=42, N2=49 |
| EE, Cluster 1: Pre/Post response; Synchrony | Figure 6K | independent t-test | t=10.832, p=<0.0001, N1=153, N2=98 |
| EE, Cluster 2: Pre/Post response; Synchrony | Figure 6K | independent t-test | t=0.921, p=0.3602, N1=19, N2=13 |
| EE, Cluster 3: Pre/Post response; Synchrony | Figure 6K | independent t-test | t=1.331, p=0.1916, N1=3, N2=12 |
| Cell counts: Restraint*Condition; Week 1 | Supp. Figure 8A | chi-square test | X^2^(1)=1.256, p=0.2624 |
| Cell counts: Restraint*Condition; Week 4 | Supp. Figure 8A | chi-square test | X^2^(1)=0.05, p=0.8223 |
| Cell counts: Restraint*Condition; Week 1-Cluster 1 | Supp. Figure 8B | chi-square test | X^2^(1)=0.015, p=0.9011 |
| Cell counts: Restraint*Condition; Week 1-Cluster 2 | Supp. Figure 8B | chi-square test | X^2^(1)=0.371, p=0.5423 |
| Cell counts: Restraint*Condition; Week 1-Cluster 3 | Supp. Figure 8B | chi-square test | X^2^(1)=0.031, p=0.8604 |
| Cell counts: Restraint*Condition; Week 1-Cluster 1 | Supp. Figure 8B | chi-square test | X^2^(1)=1.323, p=0.2501 |
| Cell counts: Restraint*Condition; Week 1-Cluster 2 | Supp. Figure 8B | chi-square test | X^2^(1)=2.257, p=0.133 |
| Cell counts: Restraint*Condition; Week 1-Cluster 3 | Supp. Figure 8B | chi-square test | X^2^(1)=2.145, p=0.143 |
| Synchrony: Intra v Inter, CUS - Cluster 1 | Supp. Figure 8C | one sample ttest, bonferroni | t=8.951, p=<0.0001, N=608 |
| Synchrony: Intra v Inter, CUS - Cluster 2 | Supp. Figure 8C | one sample ttest, bonferroni | t=5.358, p=<0.0001, N=435 |
| Synchrony: Intra v Inter, CUS - Cluster 3 | Supp. Figure 8C | one sample ttest, bonferroni | t=5.169, p=<0.0001, N=193 |
| Synchrony: Intra v Inter, EE - Cluster 1 | Supp. Figure 8C | one sample ttest, bonferroni | t=8.726, p=<0.0001, N=534 |
| Synchrony: Intra v Inter, EE - Cluster 2 | Supp. Figure 8C | one sample ttest, bonferroni | t=3.139, p=0.015, N=70 |
| Synchrony: Intra v Inter, EE - Cluster 3 | Supp. Figure 8C | one sample ttest, bonferroni | t=3.232, p=0.0206, N=26 |

**Stats Table 7.** Highlighted rows: p < 0.05
